# A linear response theory based method for prediction of large scale protein conformational changes upon ligand binding

**DOI:** 10.1101/2021.03.09.434598

**Authors:** Rajat Punia, Gaurav Goel

## Abstract

Prediction of ligand-induced protein conformational transitions is a challenging task due to a large and rugged conformational space, and limited knowledge of probable direction(s) of structure change. These transitions can involve a large scale, global (at the level of entire protein molecule) structural change and occur on a timescale of milliseconds to seconds, rendering application of conventional molecular dynamics simulations prohibitive even for small proteins. We have developed a computational protocol to efficiently and accurately predict these ligand-induced structure transitions solely from the knowledge of protein apo structure and ligand binding site. Our method involves a series of small scale conformational change steps, where at each step linear response theory is used to predict the direction of small scale global response to ligand binding in the protein conformational space (*d*_LRT_) followed by construction of a linear combination of slow (low frequency) normal modes (calculated for the structure from the previous step) that best overlaps with *d*_LRT_. Protein structure is evolved along this direction using molecular dynamics with excited normal modes (MDeNM) wherein excitation energy along each normal mode is determined by excitation temperature, mode frequency, and its overlap with *d*_LRT_. We show that excitation temperature (ΔT) is a very important parameter that allows limiting the extent of structural change in any one step and develop a protocol for automated determination of its optimal value at each step. We have tested our protocol for three protein–ligand systems, namely, adenylate Kinase – di(adenosine-5’)pentaphosphate, ribose binding protein – *β*-D-ribopyranose, and DNA *β*-glucosyltransferase – uridine-5’-diphosphate, that incorporate important differences in type and range of structural changes upon ligand binding. We obtain very accurate prediction for not only the structure of final protein–ligand complex (holo-structure) having a large scale conformational change, but also for biologically relevant intermediates between the apo and the holo structures. Moreover, most relevant set of normal modes for conformational change at each step are an output from our method, which can be used as collective variables for determination of free energy barriers and transition timescales along the identified pathway.

## Introduction

Conformational transitions are central to the biological functioning of proteins.^1,2^ Recent studies have suggested that such transitions are encoded in the protein fold and are triggered by external stimuli.^3,4^ In particular, ligand-induced conformational transitions play an important role in several biological pathways, viz. signalling,^5^ membrane transport,^6^ protein synthesis,^7^ enzyme catalysis,^8^ active transport by motor proteins,^9^ and allosteric regulation,^10^ making knowledge of protein structural response upon ligand binding very important for firmly establishing structure-function relationships.^11^ Further, prediction of ligand-induced protein structural transitions are very important for structure-based drug discovery approaches.^12^ For example, accounting for receptor conformational flexibility has been shown to improve prediction of both the structure and the binding affinity of the protein-ligand complex.^2,12,13^

Advancement in computer hardware and simulation algorithms coupled with significant progress in experimental techniques for structure determination has provided several insights into a dynamic nature of protein structure – function relationships spanning from local (at binding site) to global (at the level of entire protein molecule to protein assemblies). However, prediction of large-scale conformational transitions of proteins including identification of functionally relevant intermediates (on-pathway), either experimentally or computationally, still presents a formidable task (for example, see recent review articles on the subject^11,12^). These structural transitions span at least two orders of lengthscale (from atomic, ~ 1Å, to entire protein molecule, ~ 100Å) and typically involve large barriers on a rugged free energy landscape (FEL), leading to transition timescales of milliseconds to seconds and conformations trapped in a local minima. All atomistic molecular dynamics (MD) simulations based on the system Hamiltonian, although ideally suited to probe details on the required atomic lengthscale, are made intractable by large range of timescales involved. Several methodologies proposed to overcome this timescale problem, such as reaction path methods and potential smoothing approaches, have been shown to generate atomically detailed functional trajectories of protein transitions that provide information on structure of biologically relevant intermediates and end states as well as free energy barriers and transition timescales.^14^ These techniques, referred to as low-throughput MD methods, are still computationally intensive and this resource requirement typically increases exponentially with sum total of relevant degrees of freedom for protein dynamics.^15,16^

An alternate approach involves the use of coarsegrained dynamics in the subspace of protein degrees of freedom, with the normal modes analysis (NMA) being the most prominent. Numerous studies have established that protein structural transitions upon ligand binding can be described by a small set of slow normal modes of protein fluctuations in its unbound (apo) state.^3,4,17^-20 Emergence of highly simplified elastic network models (ENMs) to obtain these normal modes21 from a single structure, for example a crystallography structure, led to widespread use of NMA in flexible docking, targeted conformational search, and prediction of functional motions of proteins. Several enhanced sampling techniques that accelerate protein dynamics along these slow modes combined with use of coarsegrained experimental information on the structure of protein-ligand complex as constraints have shown success in predicting the holo (ligand-bound) state of protein from its apo (ligand-unbound) state.^22^ For example, structural displacement along a linear combination of low-frequency modes of the apo state, where coefficients of linear combination were determined using a small set of distance constraints of the end state, was used to predict the structure of the end state.^23^ Perahia’s group developed an enhanced sampling method, MDeNM (molecular dynamics with excited normal modes), in which protein is kinetically excited along the selected subset of modes.^24^ Using this method, end conformations are obtained with the prior knowledge of dominant normal modes involved in conformational transitions upon ligand binding. Since overall protein structure change can be non-linear, whereas only linear structure change is possible by displacement along a given set of normal modes, techniques based on small displacement driven along the predicted direction and a re-calculation of low frequency motion from the obtained structure are expected to be better suited for determination of large-scale structural changes. Tama et al. used the linear combination of slow modes to iteratively deform the structure towards target guided by low resolution electron density map of target structure.^25^ Wu and co-workers has accurately predicted the folded state of GB1 starting from its partially folded state using cryo-electron microscopy map restrained self-guided Langevin dynamics.^26,27^ All the above approaches use NMA based enhanced sampling methods guided by either low resolution structural information on end state or prior knowledge of transition pathway.

Oftentimes prior knowledge on end state or reaction coordinate is not available, for example in virtual drug screening, where incorporation of receptor flexibility notably improves screening accuracy.^13^ One option involves displacement along a predefined number of lowest frequency normal modes combined by energy minimization of the docked ligand-protein complex to predict holo state solely from its apo state^28,29^. Computational cost of these methods increases geometrically (2^*x*^) with the number of modes (*x*) selected for optimization. Moreover, inclusion of irrelevant modes would add noise in the energy calculations performed to identify the most stable ligand-bound state^30^. Therefore, to achieve high accuracy and computational efficiency, the most important choice in protein holo structure prediction methods is the selection of the relevant degrees of freedom^12^. Cavasotto and co-workers introduced a heuristic measure to determine relevance of normal modes, and thus enable exclusion of irrelevant modes^30^. This involved generation of multiple receptor conformers by perturbing structure along random linear combinations of selected relevant modes, followed by docking and energy optimization to identify the most stable ligand-bound state^30^. Another option is to identify the specific set of normal modes from the knowledge of intrinsic dynamics of protein in free state and perturbation caused by ligand binding^3,31^. Few recent studies attempted to model this predisposition of conformational transitions without any knowledge on end conformation. Zheng and Doniach predicted the conformational change in terms of Hessian matrix of protein in its unbound state and ligand induced structure distortions in binding pocket^9^. Ikeguchi and co-workers have shown that classical linear response theory (LRT) can predict the coupling relation between ligand-induced force on protein and protein conformational response^32^. Notably, the governing equations derived by Zheng and Doniach for predicting direction of conformational change is quite similar to the LRT. The direction of conformational change predicted showed a high correlation with observed direction of conformation change. However, prediction of final state using these equations is not straightforward because protein conformational change in general is non-linear and these equations do not accurately predict the extent of conformational change. Tamura and Hayashi have shown that sole application of LRT failed to accurately predict the conformational change in N-calmodulin which undergoes large scale non-linear conformational transitions upon ligand binding.^16^. Moreover, they have shown that the direction of protein response changes along the transition path and LRT can be used in a sequential manner to identify the relevant coordinate of conformational change in each new structure identified along the transition path.

Apart from the prediction of protein holo state from its apo state, the information on apo → holo transition pathway and short-lived intermediate states is crucial for understanding the biological functioning mechanisms of proteins and developing rational intervention strategies^33^. Accurate prediction of transition pathway requires a converged sampling of free energy landscape along the CVs describing the transition path which is non-trivial and computationally expensive. However, several coarse-grained morphing methods have been proposed for generation of transition pathways connecting apo and holo states. MolMovDB^34^, which performs linear interpolation between apo and holo state in Cartesian space; NOMAD-Ref^35^, which uses ENM-based normal modes to interpolate interresidue distances between specified end states; MinAction-Path^36^, which solves action-minimization problem to find the crossing point of trajectory solutions generated from the two end states; NMSim^37^, which performs biased backbone motion in slow-modes subspace followed by favourable rotamer state search for side-chains; and iMODS^38^, which performs geometric interpolation on dihedral angle space, are some of the coarse-grained methods available for use via web-servers. Recently, Orellana and co-workers have shown the application of hybrid elastic-network Brownian dynamics simulations (eBDIMS) for generating accurate trajectories connecting apo and holo states39. Although these coarse-grained methods predict the transition trajectory in a computationally inexpensive manner, the knowledge of both apo and holo high-resolution structures is required a priori.

We developed a computational protocol to efficiently and accurately predict the apo → holo transition pathway, i.e., both the intermediates and holo state, solely from the apo state of a protein. This protocol is based upon linear response theory which predicts the response of a statistical mechanical system due to weak perturbations in its Hamiltonian^32^. Using LRT, the protocol predicts the near-equilibrium response of protein structure due to the perturbation caused by ligand binding and identifies the dominant normal modes triggered by it. After predicting the direction of conformational change, it performs an enhanced conformational search along the predicted direction to identify metastable intermediates. As protein transition pathways are typically non-linear, this protocol performs an iterative prediction of response and subsequent structure transitions to reach the end state. Although, ligand binding is a non-weak perturbation to the Hamiltonian of protein, we have also derived that linear response theory will still be valid and accurately predict the protein structural response. We have successfully tested the applicability of our protocol to three protein–ligand systems, namely, adenylate Kinase (ADK) – di(adenosine-5’)pentaphosphate, ribose binding protein (RBP) – *β* -D-ribopyranose, and DNA *β* -glucosyltransferase (BGT) – uridine-5’-diphosphate. These three systems incorporates variations in scale and nature of structural transitions upon ligand binding. For example, in ADK, a sequential hinge-bending closure of its two domains is observed whereas, in RBP, a large-scale simultaneous twisting and hinge-bending motion of two of its domains is observed. Also, the RMSD between apo and holo states varies from 2.1 Å (for BGT) to 7.1 Å (for ADK).

## Theory

For a statistical mechanical system in equilibrium, LRT predicts its first order response to perturbation in its Hamiltonian^40^. For a protein-ligand system, ligand binding perturbs the Hamiltonian of unbound-protein and protein change its state to reach the new free energy minima. The Hamiltonian of ligand-bound state 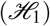 can be written in terms of the Hamiltonian of ligand-unbound state 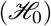 as

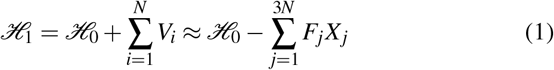

*V_i_* is the interaction energy between ligand and *i^th^* residue of protein, *N* is the total number of residues, *X_j_*s are the generalized coordinates such that {*X*_1_,*X*_2_,...*X*_3*N*_} = {*x*_1_,*y*_1_,...*z_N_*} and Fs are the forces exerted by the ligand on residues such that {*F*_1_, *F*_2_,... *F*_3*N*_ } = {*f*_*x*_1__, *f*_*y*_1__,... *f_z_N__* }, where {*x*_1_, *y*_1_, *z_i_*} are coordinates of *i^th^* residue and *f*_*x*_i__, *f*_*y*_i__ and *f*_*z*_i__ are the ligand-exerted forces on *i^th^* residue in x,y and z direction. The first order response of protein structure due to ligand-induced perturbation given by the LRT-formula (equation A11) is^32^

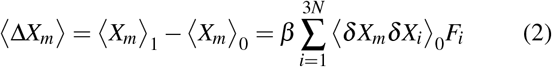

〈·〉_1_ and 〈·〉_1_ denotes ensemble averages in states before and after perturbation, *β* (= 1/*k_B_T*) is thermodynamic beta and 〈*δX_i_δX_j_*〉_0_ is covariance between *X_i_*, and *X_j_*. LRT-predicted structural response is function of fluctuation covariances of protein residues and the perturbative forces exerted by ligand (equation 2), and both the properties depends upon the conformational state of the protein. In other words, the LRT-predicted response changes as the protein conformational state changes. Generally, in large-scale conformational changes upon ligand binding, protein observes transitions from initial to final state via multiple intermediate states^2^, and, in general, the response will change along the transition coordinate resulting in a non-linear transition pathway^16^ (discussed later). Therefore, it would be appropriate to use equation 2 only for the prediction of local response of protein motion from a given state to the adjacent state along the transition path but not for the prediction of the complete response due to ligand binding. Moreover, although a good correlation exists between the LRT-predicted direction of conformational change and actual direction conformational change, the magnitude or extent of conformational change is typically incorrectly estimated by the LRT-formula (Supplementary Table S1). Therefore, in our protocol we use LRT-formula to predict only the direction (not magnitude) of the structural response (**d**_*LRT*_) and consequently the intrinsic motions of protein in a given state triggered by ligand binding.

To identify the dominant normal modes triggered by ligand binding in a given conformational state (S), we first predicted the direction of conformational change (**d**_*LRT*_) using LRT in state S and then calculate the overlap (*O_i_*) of LRT-predicted direction with each normal mode (**U_i_**) of state S. Overlap with *i^th^* mode is calculated as

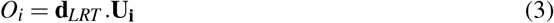

Normal modes are mutually orthogonal and **d**_*LRT*_ can be represented by linear combination of normal modes as

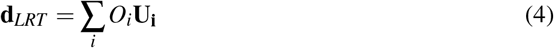

Modes having a large overlap are the dominant modes triggered by the ligand binding. These dominant modes can possibly be used as CVs in enhanced sampling methods to describe the ligand-induced structural response of protein.

Besides, as LRT predicts the first order response which is accurate for weak perturbation in Hamiltonian, the question arises on the accuracy of LRT-predicted direction as ligandbinding can cause strong perturbation in protein Hamiltonian. Therefore, to examine the validity of LRT to predict direction of conformational change, we derived *(SI Index)* the general response (equation 5) of protein structure due to ligand-induced perturbation in protein Hamiltonian which includes higher order terms.

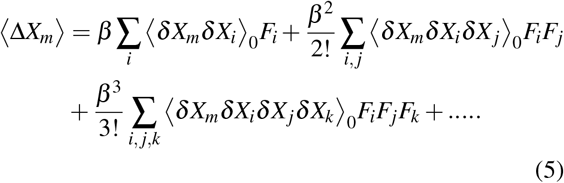

Assuming a harmonic approximation of potential energy of protein, which is valid for small fluctuations near minima of an energy well, residues displacement from its mean position follows a multivariate Gaussian distribution^41^. For a multivariate Gaussian distribution of residue positions, we derived that (*SI Index*), each higher order term in the equation 5 is proportional to the first term and the general response is given as

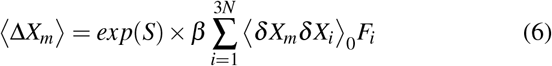

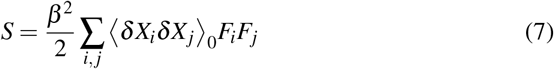

Equation 6 shows that the general response of protein is proportional to the linear response. Therefore, as far as the direction, and not the magnitude, of initial structural response from a given state to an adjacent state is concerned, it would be appropriate to use LRT for strong ligand-induced perturbations.

## Results

Application of our protocol to predict protein structure transitions upon ligand binding is shown for three protein+ligand systems: (i) Adenylate Kinase (AdK) + Di(adenosine-5’)pentaphosphate(Ap5A) (ii) Ribose Binding Protein (RBP) + β-D-ribopyranose (RIP) and (iii) DNA β - glucosyltransferase (BGT) + Uridine-5’-Diphosphate (UDP). The main steps followed in our protocol are summarized below:

1. Docking of ligand on protein near the binding site followed by solvation, equilibrations and 1 ns long explicit-solvent MD relaxation (see Methods).
2. Estimation of ligand-induced force on protein residues and prediction of direction of protein structural response using LRT.
3. Identification of dominant normal modes triggered by ligand binding and enhanced sampling along LRT-predicted direction using MDeNM.
4. Identification of closest distinct conformational state along the excited direction and repetition of steps (2)-(4) (multiple iterations)
5. Identification of most-stable ligand-bound state (holo state) among the output states obtained from multiple iterations

### Identification of intrinsic motions triggered by ligand binding

We performed normal mode analysis (NMA) of protein fluctuations to identify the intrinsic motions of protein. NMA decompose the thermal fluctuations of protein into a set of orthogonal modes of vibration called as normal modes. We calculated normal modes by modelling protein as anisotropic network model (ANM)^42^, in which each protein residue is modelled as a single bead centred at its Cα atom and each bead is connected to other beads within the cutoff radius *(R_c_*) by harmonic springs of uniform spring constant (*γ*). The potential energy function *(E)* of this coarse-grained representation of protein is

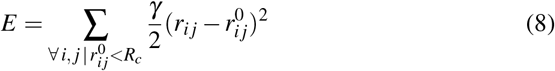

*r_ij_* and 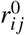 are the instantaneous and equilibrium distance between residues *i* and *j*. By carefully selecting the two parameters (*R_c_, γ*), this elastic network model, accurately and efficiently predict slow and collective modes of protein fluctuations^42^. Relative fluctuations of residues depends upon the value of *R_c_*, while *γ* only changes the overall scale of fluctuations. Thus, most important ANM parameter is *R_c_*, which we chose on the basis of two factors:

#### (i) ANM should accurately predict the thermal fluctuations of protein

For this, we calculated the Pearson correlation coefficient *(ρ*) between residue fluctuations obtained from MD simulations and estimated using ANM for different values of *R_c_* (see Methods). The maximum value of *ρ* obtained for AdK, RBP and BGT is 0.91, 0.80 and 0.78 respectively (Fig. 1, Supplementary Fig. S1, S2). High value of *ρ* for our test cases implies good prediction of thermal fluctuations by ANM, thus allows its use for normal mode calculations. For a given protein, we selected a set comprising all *R_c_* values which gives correlation greater than 95 percent of the maximum correlation. The selected set of *R_c_* values for AdK, RBP and BGT are [8 — 13 Å], [8 — 10 Å] and [8 — 10,12 Å] respectively (Fig. 1, Supplementary Fig. S1, S2). The selected set contains *R_c_* values of ANM which accurately predicts thermal fluctuations of protein and from this set, a single value is selected on the basis of second factor.

**Figure 1.**
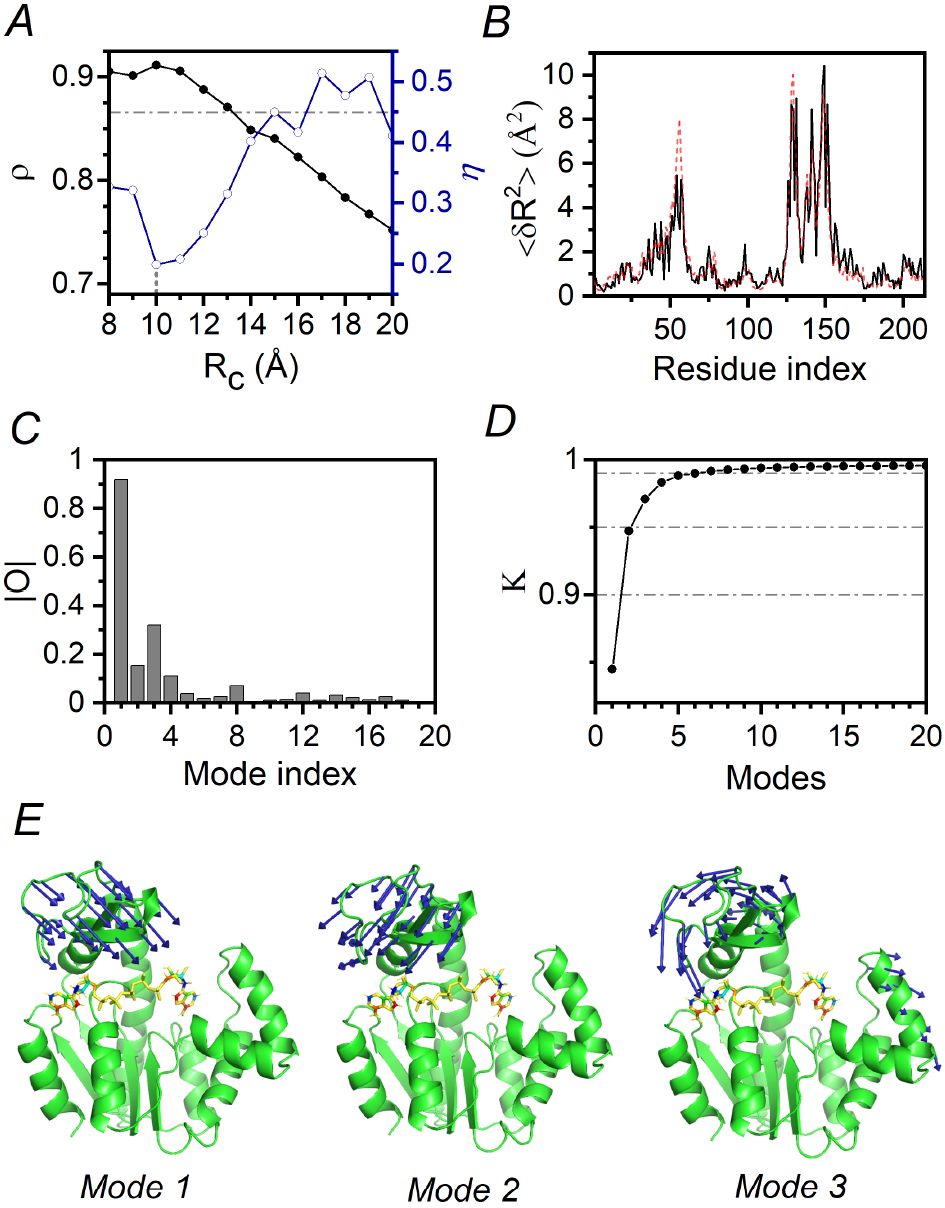
Estimation of ANM parameters and identification of normal modes triggered by ligand binding (Adenylate Kinase). (**A**) Pearson correlation (*ρ*) between residue mean squared fluctuations predicted by ANM and estimated by 1 ns long MD simulations at different *R_c_* values (filled circles, left Y-axis). A cutoff line at *ρ* = 0.95 × *ρ_max_* is shown (dashed line). Mode contribution diversity estimated by *η* for different *R_c_* values (empty circles, right Y-axis). (**B**) Mean squared fluctuations of residues in apo-state obtained from 1ns MD simulation (dotted line) and estimated by ANM at *R_c_ =* 10Å (solid line). (**C**) Overlap (absolute value) of first 20 normal modes with LRT-predicted direction. (**D**) Cumulative contribution of top normal modes to LRT-predicted direction in decreasing order of overlap (circles) and reference lines at *K* = 0.9, 0.95 and 0.99 (dotted lines). (**E**) Direction of motion of residues in *Mode* 1, *Mode* 2 and *Mode* 3 shown by blue arrows on cartoon diagram of protein.

#### (ii) the identified set of relevant CVs describing the predicted direction of conformational change should be small

For each values of *R_c_* from the selected set, we obtained the normal modes and their overlap with the LRT-predicted direction (**d**_*LRT*_). As discussed in theory section, modes having large overlap are the dominant modes triggered by ligand binding. These dominant modes could be selected as CVs to perform enhanced sampling and it is desirable for optimal performance of enhanced sampling methods that the set of relevant CVs should be small. In other words, since **d**_*LRT*_ can be represented by linear combination of normal modes (equation 4), it is desirable that most of the contribution to **d**_*LRT*_ is distributed among a small set of normal modes. We defined a Shannon-entropy based measure *η* to quantify the diversity of contribution of first *M* normal modes to **d**_*LRT*_.

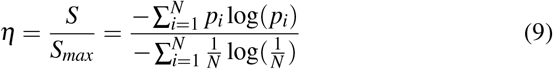

Here, 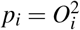 as 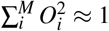. The value of *η* lies between 0 and 1 for extreme cases. The case *η* = 1 means all *M* modes contribute equally to **d**_*LRT*_ and case *η* = 0 means **d**_*LRT*_ is completely represented by a single mode. Thus, from the selected set of *R_c_* values, we select a single *R_c_* which corresponds to minimum value of *η* (Fig. 1, Supplementary Fig. S1, S2). Lower value of η implies most of the contribution to **d**_*LRT*_ comes from a small set of modes. Selected *R_c_* value on the basis of these two factors ensures that ANM accurately estimates the intrinsic fluctuations of protein residues as well as structural response (**d**_*LRT*_) can be described by a small set of collective modes.

Normal modes obtained from final selected *R_c_* value are used for further analysis. Overlap of first 20 modes with the LRT-predicted initial response is shown for AdK, RBP and BGT (Fig. 1, Supplementary Fig. S1, S2). For all test case, few slowest frequency modes have the high overlap (relevant modes) with LRT-predicted direction, whereas rest of the modes contribute very low (non-relevant modes). In order to select only relevant modes, we defined a parameter (*K*) to measure the cumulative contribution of top *M* normal modes. Value of *K* is zero for *M* = 0 and approaches 1 for all normal modes.

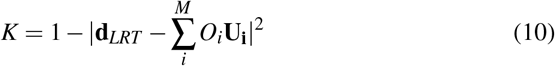

Summation in equation 10 is over most dominant *M* modes in decreasing order of overlap. Smallest set of top modes whose cumulative contribution is greater than 95% is designated as the set of relevant modes triggered by ligand binding (Fig. 1(D)). We tested 3 cutoff values 90%, 95% and 99% and observed the best structure match of predicted and experimental holo-structure with 95% cutoff (data not shown). A lower cutoff value (90 %) clearly exclude some of the relevant modes, while a higher cutoff (99 %) value includes many low-overlap modes which adds noise to the calculations. Initially, 3, 3 and 2 normal modes are triggered by ligand binding in AdK, RBP and BGT respectively (Fig. 1, Supplementary Fig. S1, S2). These relevant modes are the intrinsic motions triggered by ligand binding which can potentially be used as CVs in enhanced sampling methods simulating ligand-induced protein structure transitions. We use the selected relevant modes to perform sampling along the LRT-predicted direction using MDeNM, a recently developed computationally efficient enhanced simulation method.

### Enhanced sampling along LRT-predicted direction of conformational change using MDeNM

Enhanced sampling along the LRT-predicted direction of conformational change is performed using MdeNM, which is modified to perform a targeted sampling along the LRT-predicted direction. In typical MdeNM simulations, extra velocity (λ**v**_*extra*_) is added to the current velocity (**v**_0_) to excite protein structure along selected modes^24^.

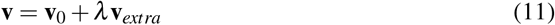

Here, **v** is the updated velocity of protein, **v**_*extra*_ is linear combination of selected modes (**v**_*extra*_ = ∑_*i*_ *α_i_***U_i_**; *α_i_* is velocity coefficient of *i^th^* mode) and *λ* controls the degree of excitation such that the temperature increase of system (Δ*T*) due to velocity excitation is

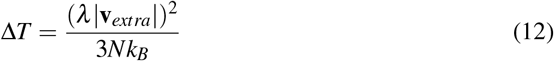

*k_B_* is Boltzmann constant and *N* is number of atoms in the system^24^. Costa et. al. discussed the efficacy of MdeNM for random sampling (**v***extra* is random linear combination of selected modes) and targeted sampling along a single mode^24^.

Generally, the LRT-predicted direction is represented by linear combination of multiple normal modes. Thus, an effective strategy is required to perform a targeted sampling along LRT-predicted using MDeNM. To perform, a targeted sampling along a pre-determined direction, velocity coefficients of normal modes need to be carefully decided. Taking velocity coefficients proportional to the predicted overlap *(i.e. α_i_* ∞ *O_i_*) doesn’t sample the conformations in targeted direction because of unequal stiffness of modes. For example, a unit displacement along a low-frequency mode requires less excitation velocity compared to a high-frequency mode or, a given excitation velocity along the slow mode will produce a larger conformational change compared to the excitation with same velocity along a fast mode (Supplementary Fig. S3). Assuming harmonic fluctuations along a given mode i, the maximum displacement along the mode (*d_max,i_*) is governed by the energy balance equation as

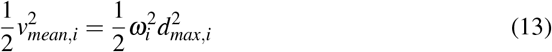

*ω_i_* is frequency of *i^th^* mode and *v_mean,i_* is velocity at equilibrium position along *i^th^* mode. Thus, based on the harmonic approximation of protein fluctuations along the normal modes, near an energy minima, we decided velocity coefficient of *i^th^* mode to be proportional to the predicted overlap along the *i^th^* mode (*O_i_*) and mode frequency *ω_i_* (*i.e. α_i_* ∞ *ω_i_O_i_*).

MDeNM kinetically excite the protein motion along a predetermined direction. This excitation is equivalent to impulse, in which protein velocities are suddenly increased along the excited direction thus increasing the temperature of the system which is quickly reduced to set temperature by thermostat in simulation (Supplementary Fig. S3). Although, the extra kinetic energy is quickly dissipated by thermostat, the kinetic energy along the excited direction remains high relative to other degrees of freedom (DOF) because thermostat extract extra kinetic energy from all DOF of the system. Thus, the protein displacement along the predicted direction continues until the kinetic energy gets partitioned equally (on an average) among all DOF of the system. Therefore, in a MDeNM excitation, the initial response of protein structure is governed by the excitation temperature (Δ*T*) which is quickly dissipated by thermostat, subsequent response is governed by the protein’s Hamiltonian or the underlying free energy landscape and total structure response is governed by both Δ*T* and protein’s Hamiltonian. Costa et. al. discussed in detail, how protein structure changes during a MDeNM excitation.^24^ Briefly, upon excitation, *D_P_* increases suddenly, governed by the value Δ*T*, reaching a maximum value 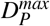 within ~ 5 *ps*, followed by relaxation of structure (and *D_P_*) towards the nearest energy minimum state (Supplementary Fig. S3).^24^ More will be discussed on the characteristics of initial and total structure response with varying Δ*T* in the next section.

### Identification and validation of protein holo-state and transition intermediates

RMSD between the *C_α_* atoms of apo and holo crystal structures of AdK, RBP and BGT is 7.1, 6.2 and 2.1 Å respectively. After 1ns MD relaxation of docked structure (state *I*_1_), large structure deviation remains. RMSD between *C_α_* atoms of *I*_1_ and holo cryatal structures of AdK, RBP and BGT is 6.1, 5.2 and 3.7 Å respectively. During this small MD simulation, protein-ligand complex relax to the nearest free energy minima. Therefore, for a given protein, large perturbation in the protein structure is required to reach near-holo states, which are usually separated by large free energy barriers. Large perturbation in initial structure can be carried out using two approaches (i) a single large perturbation along the LRT-predicted direction or (ii) multiple small perturbations along the LRT-predicted directions. Importantly, the second approach allows updated calculation of direction of conformational change before each stage of small perturbation.

First approach performs a single large pertrurbation of initial structure and, in MdeNM, the extent of protein structure change along the predicted direction (measured as *D_P_* = root mean square displacement of residues along excited direction; see Methods) is controlled by changing the value of excitation temperature (Δ*T*). However, as discussed in theory section, *D_P_* cannot be accurately estimated using LRT and thus, deciding the value of Δ*T* in advance is not possible. Therefore, we observed the protein structure change at varying degree of excitation, ranging Δ*T* from 0 K to 200 K at the gap of 5 K. Since, the initial response of protein in a MDeNM excitation is governed by Δ*T*, we observed a continuous increase in 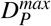 with increasing Δ*T*. Whereas, the final response (after MD relaxation) does not increase continuously with Δ*T* and a three-point average shows the presence of several plateaus in *D_P_* vs Δ*T* plot (Fig. 2, Supplementary Fig. S4). Plateaus corresponds to an approximately same *D_P_* for a range of Δ*T*, which indicates the presence of stable region along the transition coordinate **d**_*LRT*_. Structures obtained in the range of Δ*T* between two consecutive plateaus indicates the unstable region along the transition coordinate *D_P_*. Structures obtained in these unstable regions probably corresponds to states trapped in the local energy minimas in rugged energy landscape of the protein. Output structures obtained at the end of each excitation are compared with the experimental holo structure (Fig. 2, Supplementary Fig. S4) by calculating *RMSD* and *GDT_avg_* score (see Methods). Protein structure improvement, estimated as decrease in RMSD with the experimental holo structure after excitation, has strong correlation (0.91, 0.84 and 0.96 for ADK, RBP and BGT) with *D_P_* till the end of first plateau. The correlation starts to decrease for higher Δ*T* values and even becomes negative for very high Δ*T* values (Supplementary Table S2). Since protein structure improvement is strongly correlated with displacement along LRT-predicted direction, we assigned representative structures (*P*_1_, *P*_2_,...) to every plateau corresponding to highest *D_P_* value in each plateau region (see Methods). Our protocol aims to predict the holo state solely from the apo state, hence, it is assumed that no information of any kind is available on holo state. Holo state is the energetically most stable ligand-bound state of a protein, thus, we identify the most-stable representative structure using a energy-based scoring function (ΔΔ*G*) and designate it as the predicted holo state. ΔΔ*G* accounts for changes in protein-ligand binding energy and protein strain energy due to protein structure changes (see Methods). Using first approach, the RMSD between predicted holo structures and crystallographic holo structures for AdK, RBP and BGT is 3.79, 3.94 and 2.36 respectively (Table 1, Supplementary Table S3, S4).

**Figure 2.**
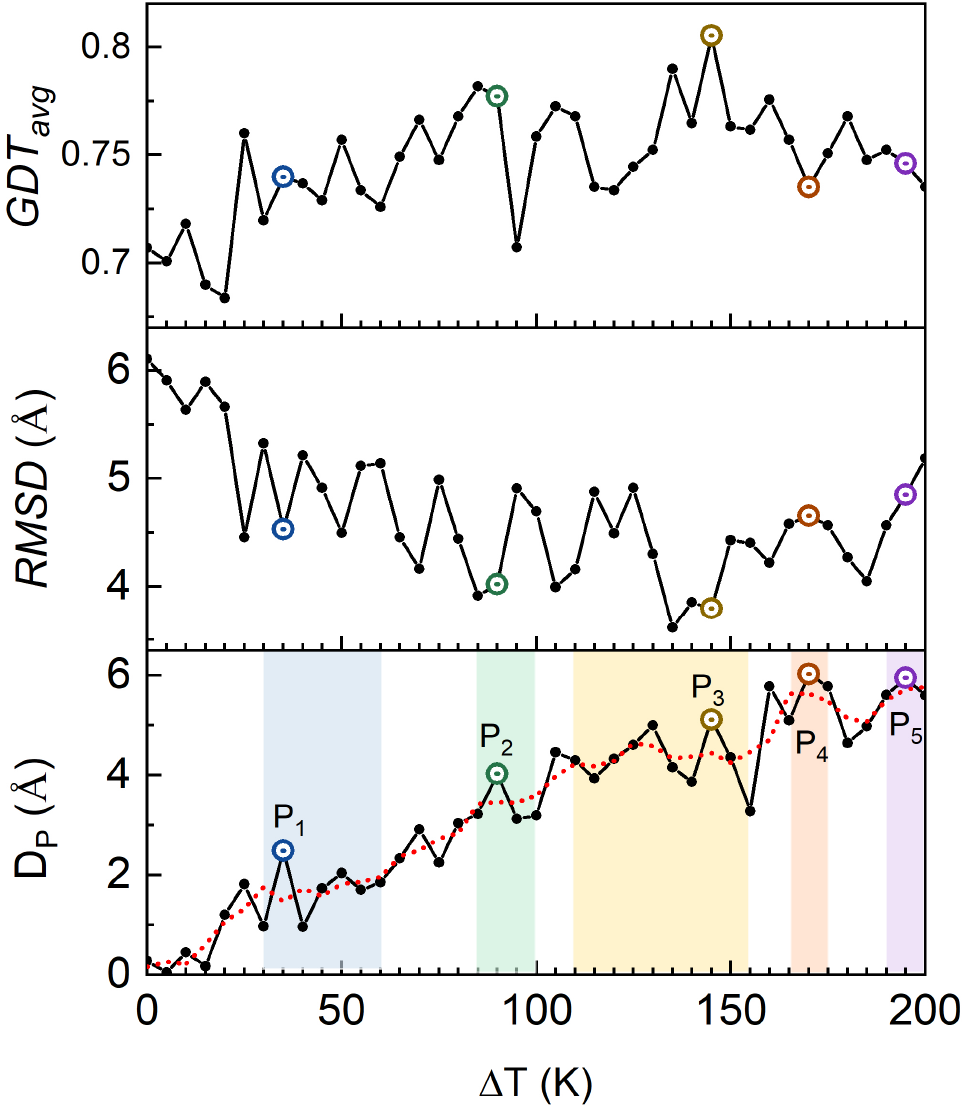
Enhanced sampling along LRT-predicted direction (Adenylate Kinase). Structure displacement along LRT-predicted direction (*D_P_*) in 100 ps MDeNM excitations at varying Δ*T* (circles, solid line) along with three-point average of *D_P_* vs. Δ*T* data (dotted line). Identified plateau regions in average *D_P_* vs. Δ*T* plot are highlighted (bottom panel). *RMSD* (middle panel) and *GDT_avg_* (top panel) score between the experimental holo structure and structure obtained at the end of MDeNM excitation. Representative structure of each plateau is highlighted (all panels).

**Table 1.** Identification of holo struture from energetics (Adenylate Kinase).

| | $RMSD (A^0)$ | $GDT_{avg}$ | $\Delta\Delta G (Kcal/mol)$ |
| --- | --- | --- | --- |
| *Initial* | 7.14 | 0.66 | - |
| *I*<sub>1</sub> | 6.10 | 0.71 | 0 |
| Single large perturbation |  |  |  |
| *P*<sub>1</sub> | 4.53 | 0.74 | 0.68 |
| *P*<sub>2</sub> | 4.02 | 0.77 | -22.46 |
| ***P*<sub>3</sub>** | **3.79** | **0.80** | **-26.04** |
| *P*<sub>4</sub> | 4.65 | 0.73 | -11.00 |
| *P*<sub>5</sub> | 4.85 | 0.75 | -5.62 |
| Multiple small perturbations |  |  |  |
| *E*<sub>1</sub> | 4.53 | 0.74 | 0.68 |
| *E*<sub>2</sub> | 4.01 | 0.77 | -0.66 |
| *E*<sub>3</sub> | 3.78 | 0.78 | -12.75 |
| *E*<sub>4</sub> | 2.72 | 0.85 | -29.68 |
| ***E*<sub>5</sub>** | **2.30** | **0.87** | **-61.49** |
| *E*<sub>6</sub> | 2.49 | 0.85 | -52.37 |
| *E*<sub>7</sub> | 2.75 | 0.83 | -36.30 |
| *E*<sub>8</sub> | 2.76 | 0.83 | -43.42 |

*Initial* = docked protein-ligand structure
*I*<sub>1</sub> = structure after 1 ns MD relaxation of *Initial*
*P*<sub>*i*</sub> = representative structure of *i*<sup>th</sup> plateau
*E*<sub>*j*</sub> = representative of first plateau in *j*<sup>th</sup> perturbation

In the second approach, large-scale structure changes, in initial state, to reach near-holo state is achieved by multiple small perturbations along the LRT-predicted directions. As the first plateau in the *D_P_* vs Δ*T* plot corresponds to smallest significant transition along the LRT-predicted direction, we selected the representative structure of first plateau as the output to perform the next perturbation. Protein fluctuations, fluctuation covariance matrix and ligand-induced forces depends upon protein conformational state and thus change after each perturbation. Therefore, before each perturbation the direction of conformational change is recalculated with updated covariance matrix and ligand-induced forces. Multiple such iterations (~ 10 *iterations)* of prediction of conformational change and subsequent small perturbation along the predicted directions are performed. Output of each iteration is analysed using the scoring function ΔΔ*G* and the most stable output state is designated as the final state. This final state is the predicted holo-state of protein and all the states connecting the transition from inital to final state are designated as intermediate states. For all three proteins, the most stable state predicted by the scoring function corresponds to the state structurally most similar to the experimental holo state as measured by *RMSD* and *GDT_avg_* score (Table 1, Supplementary Table S3, S4). RMSD between predicted holo structures and experimental holo structures for AdK, RBP and BGT is 2.30, 1.58 and 1.4 Å respectively (Table 1, Supplementary Table S3, S4). Unsurprisingly, the holo structure predicted using the second approach have higher similarity with the experimental holo structure for all proteins. Large scale-structure changes, typically follow a non-linear pathways, in which intermediate states play an important role in governing and modulating conformational transitions. Since, the protein conformational transition pathways can be non-linear, the single large perturbation along the LRT-predicted direction is not the best approach, as it ignore the important role of transition intermediates. These valid concerns are eliminated in the second approach which performs small but multiple iterative structure perturbations, incorporating the role of intermediate states in modulating the direction of transition pathway.

Fig. 3 shows the structural comparison of predicted and actual holo structures of the three proteins. Further, we calculated and compared the displacement of the predicted and experimental holo structures from the initial structure along the first 10 normal modes of the initial structure (Fig. 3). Most of the contribution to the overall displacement comes from the first few low frequency modes. Correlation between predicted and actual displacement along the first 10 modes for ADK, RBP and BGT is 0.98, 0.97 and 0.99 respectively. Although, perturbation of initial structure along a single normal mode could improve the structure similarity with the experimental holo structure, above results shows that the high similarity between predicted and actual holo structure in our case is due to correct displacement of initial structure along multiple normal modes. Apart from the validation of predicted holo structure, we analysed whether the transition intermediates/pathway predicted by our method could recapitulate the known intermediates/pathway. We selected AdK and RBP for this analysis as the information on their transition pathway and intermediates is available from past studies.

**Figure 3.**
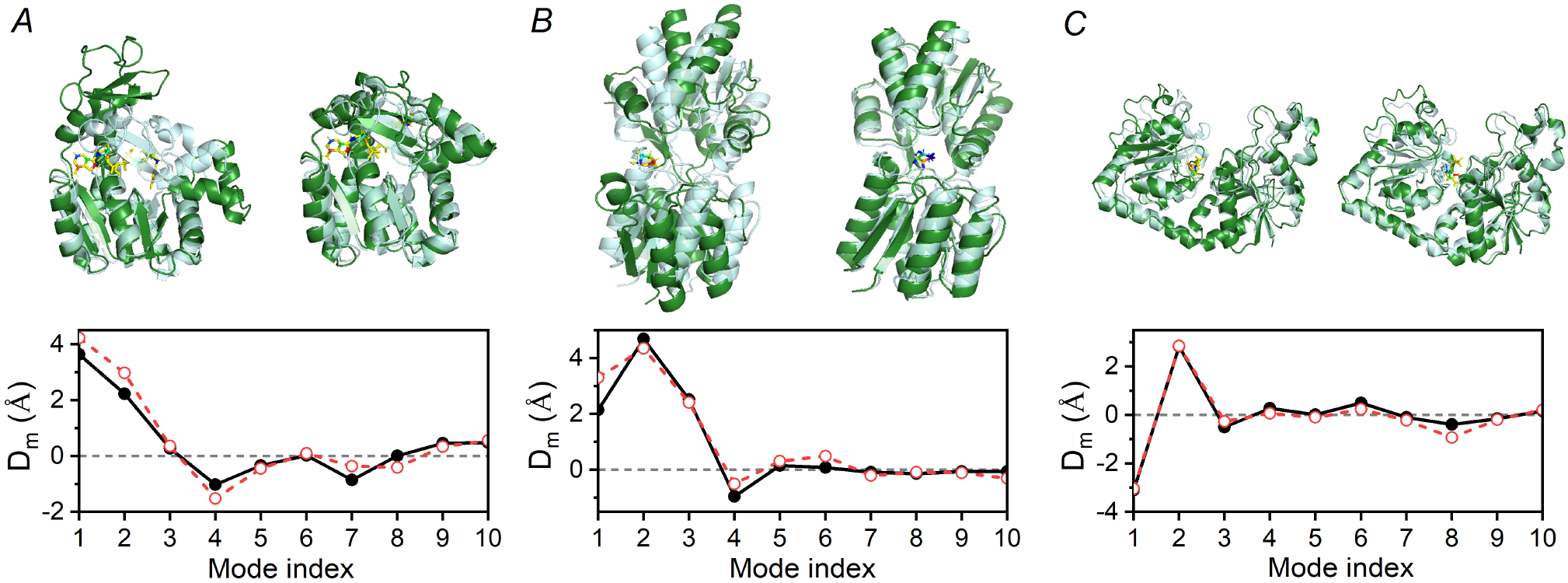
Comparison of predicted and experimental holo structures. Top panel: Cartoon diagram of initial (left) and predicted (right) holo structure (green) superimposed on experimental holo structure (Cyan); Bottom panel: Mean square displacement of predicted (black filled circles) and experimental (red empty circles) holo structure from structure *I*_1_ along first 10 normal modes of *I*_1_ for protein (A) AdK, (B) RBP and (C) BGT

#### Adenylate Kinase

AdK is an ubiquitous allosteric enzyme which catalyzes the phosphorylation reaction, *AMP* + *ATP* ↔ 2 *ADP*.^43^ This reaction involves reversible transfer of phosphoryl group from ATP (Adenosine triphosphate) to AMP (Adenosine monophosphate) which is imporatant for cellular energy homeostasis^8^. Structure of AdK comprises three domains: CORE, LID (ATP binding domain) and NMP (AMP binding domain). AdK, upon binding to ATP and AMP, undergoes an open (inactive) → closed (active) conformational transition, in which the flexible LID and NMP domains close in towards the rigid CORE domain (Fig. 4).^43^ Di(adenosine-5’)pentaphosphate(Ap5A), an inhibitor of AdK, is commonly used to simulate the combined binding effect of AMP and ATP on AdK structure.^8,43,44^ Li and co-workers computed the free energy landscape (FEL) of AP5A-bound AdK and mapped the open → closed transition pathway using bias-exchange metadynamics simulations.^8^ We projected the transition intermediates, obtained by our protocol, in sequential order on the CV space *(θ_LID_, θ_NMP_*) defined by Li and co-workers^8^. Our predicted transition pathway is consistent with their calculations, indicating, on average, the closing of LID domain first followed by the closing of NMP domain (Fig. 4). Further, we observed a state-dependent dynamic coupling between the motions of LID and NMP domains. In state *I*_1_, which is obtained after 1 *ns* MD relaxation of initial docked state, three normal modes are dominantly triggered by the ligand binding in the order *Mode* 1 > *Mode* 3 > *Mode* 2 (Fig. 1(E)). Protein internal motion in all three modes involves closing of LID domain towards the CORE domain, although along the slightly different directions. Motion in *Mode* 1 and *Mode* 2 corresponds exclusively to the LID domain closure, motion in *Mode* 3 corresponds to the coupled motion of both LID and NMP domain (Fig. 1(E). In state I_1_, motion in *Mode* 3 shows that the closure of LID domain is dynamically coupled with opening of NMP domain. Therefore, exclusive closing motion of LID domain in *Mode* 1 and *Mode* 2 and anti-correlated motion of LID closure with NMP closure in *Mode* 3 explains why, initially, NMP closing is unfavourable and LID closing is observed. In state *l*4, we observed that LID domain interacts with NMP domain and the enthalpic interactions between LID and NMP domain appears to favour the NMP closure as a large closing of NMP domain is observed thereafter, *I*_4_ to *I*_5_ (Fig. 4). In state *I*_4_, two modes are dominantly triggered by ligand binding in the order *Mode* 1 > *Mode* 2 (Supplementary Fig. S5). Both dominant modes triggered in state *1*_4_ involves the closing of NMP domain. *Mode* 2 corresponds exclusively to the closing of NMP domain, whereas *Mode* 1 corresponds to the twisting + closing of LID domain and closing of NMP domain. Interestingly, closing of LID domain is now correlated with the closing of NMP domain (Supplementary Fig. S5). The state-dependent coupling between the motions of LID and NMP domains explains why the characteristic non-linear pathway (LID closing followed by NMP closing) is observed in AdK upon ligand binding.

**Figure 4.**
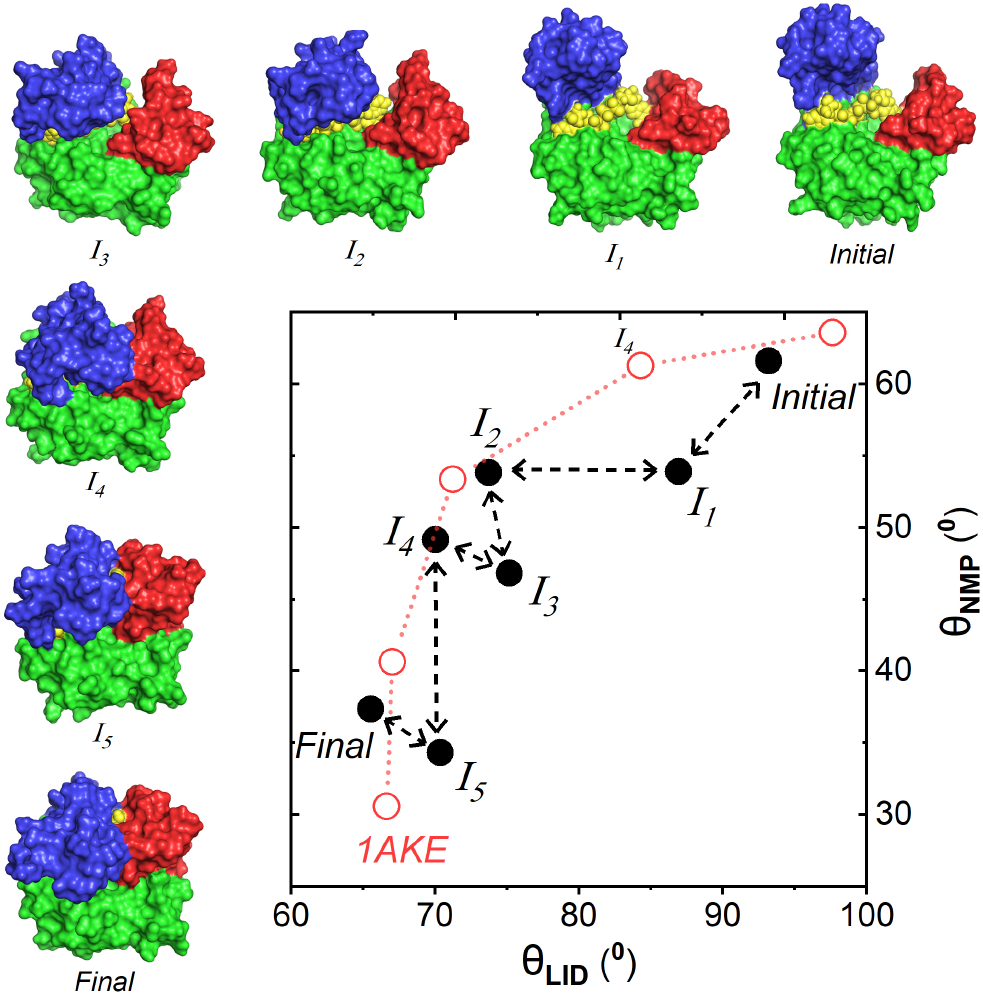
Adenylate Kinase apo → holo transition intermediates. Transition intermediates predicted by our protocol (filled black circles) along with the minimum energy path and intermediates predicted by Li and co-workers8 using bias exchange metadynamics. Surface diagram of predicted protein intermediates is also shown with colour scheme: Green (CORE domain), Blue (LID domain) and Red (NMP domain), Yellow (Ligand). Definition of collective variables *θ_LID_* and *θ_NMP_* can be found in *SI Index.*

#### Ribose Binding Protein

RBP is a bacterial periplasmic protein which mediates ribose transport and chemotaxis.^46^ It undergoes a large scale open → closed allosteric structure transition upon ribose binding, which enables its recognition by inner membrane permease responsible for ribose transport.^45,47^ Using our protocol, we also observed an open to closed transition of RBP upon ribose binding evident by decrease in protein radius of gyration by 0.82 Å. Fig. 5(A) shows the initial, final and intermediate structures predicted for RBP transition upon ribose binding. The two domains of RBP undergoes a hinge bending and twisting motion, commonly represented by order parameters hinge angle (θ) and twist angle (φ) (SI *Index).* Fortunately, the crystallographic structures of RBP provides an approximate view of the path of open → closed transition in the order, *1BA2_A → 1BA2_B* → 1URP → 2GX6 → 2DRI.^39^ To compare our transition pathway with the approximate pathway predicted from crystallographic structures, we calculated the RMSD of the structures we obtained along the transition path with the above crystal structures (Fig. 5(B)). The predicted intermediate states closely matches with the crystallographic proposed intermediates with the correct order of transitions. Intermediate *I*_1_ is closest to crystal structure 1 *BA*2_*B* with rmsd of 1.39Å, intermediate *I*_2_ is closest to crystal structure *1URP* with rmsd of 1.73 Å and final structure is closest to crystal structure 1URP with rmsd of 1.58 Å.

**Figure 5.**
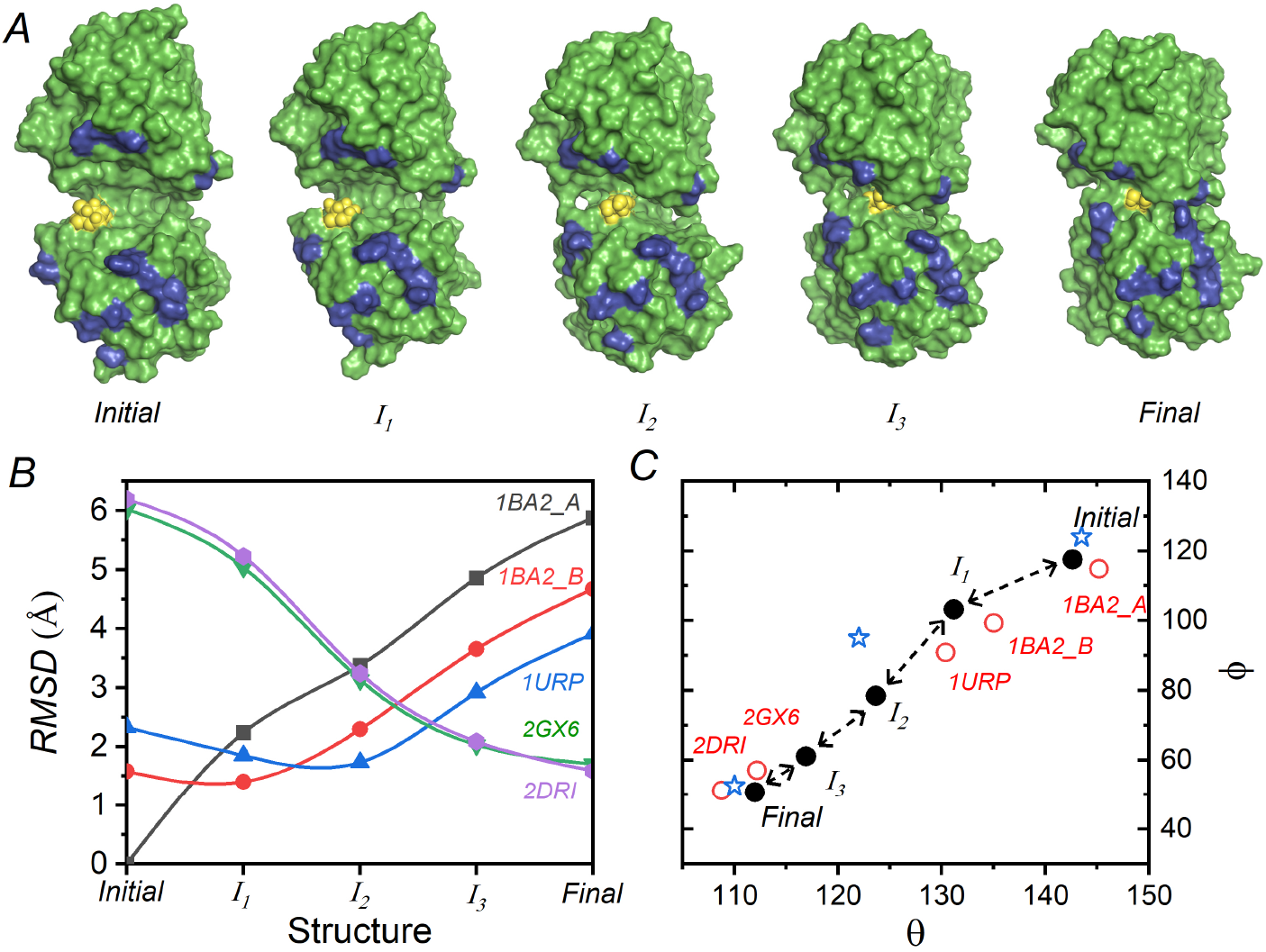
Ribose Binding Protein apo → holo transition intermediates. (**A**) Surface diagram of predicted protein intermediates. Residues which affect ribose transport but not ribose binding are coloured blue and ligand is shown as yellow spheres. (**B**) *RMSD* comparison of predicted intermediates with crystal structures. (**C**) Transition intermediates predicted by our protocol (filled circles) along with the crystallographic structures (empty circles) and metastable states of RBP predicted by Ravindranathan and co-workers^45^ using umbrella sampling (stars). Definition of collective variables *θ* and *φ* is provided in *SI Index.*

Conformational transition in RBP is important not only for ribose uptake, but also for ribose release upon interaction with inner-membrane-permease.^46^ Ribose is almost completely buried in closed state of RBP (only 20 percent of ligand surface is exposed to solvent environment) and thus it is hypothesized to undergo a partial re-opening for ribose release.^47^. The fraction of ligand surface area buried in the *Initial*, *I*_1_, *I*_2_, *I*_3_ and *Final* states is 0.47, 0.54, 0.55, 0.75 and 0.80 respectively. From the surface area analysis, it seems that state *I*_2_ is the closest (to final state) partially-open state which could release the ligand as approximately 55 percent of ligand-surface area is buried. Ravindranathan and co-workers determined the FEL of RBP conformational transition upon ribose binding using umbrella sampling MD simulations.^45^ Our predicted transition pathway is consistent with their proposed pathway, passing close to the valley regions in FEL (Fig. 5(C)). Close to the holo state, they identified a partially-open ligand-bound state (*θ* = 122, *φ* = 95) on the transition path, which they proposed to be the state relevant for ribose release. This state is closest to the state *I*_2_ (*θ* = 123, *φ* = 79) in the CV space, among all the intermediate states predicted by our protocol.

## Discussion

One underlying hypothesis, supported by large number of experimental and computational studies, that has formed the basis of method developed here is that the structural transitions of proteins often correlate with their intrinsic motions in ligand-unbound (apo) state. Therefore, ligand binding is assumed to not introduce new large-scale motions in protein, but only amplify one or more of already existing low frequency motions. Easily accessible paths or minimum energy paths are often connected with these low frequency modes because both are associated with low free energy barriers, although still much larger than thermal energy (~ *k*_B_*T*). Importantly, these collective intrinsic motions can be identified easily with coarse-grained models such as ENM without explicitly sampling the free energy landscape using an all-atomistic force field. In this work, we used LRT to identify this subset of intrinsic motions that are triggered by ligand binding and used MDeNM simulations that allow crossing of barriers on the transition pathway in a short simulation by adding kinetic energy along selected normal modes. We show that excitation temperature (Δ*T*) is a very important parameter that allows limiting the extent of structural change in any one step and develop a protocol for automated determination of its optimal value at each step.

Using the developed method, we predicted the structural response of three proteins: ADK, RBP and BGT upon binding to their ligands Ap5A, RIP and UDP respectively. We have shown that a small (~ 1 *ns*) MD relaxation after docking is not sufficient to observe the large-scale ligand-induced structure transitions. This is evident from the large RMSD between the ligand-bound structures obtained after MD relaxation of docked structure and experimental ligand-bound structures, which for the case of ADK, RBP and BGT was 6.1, 5.2 and 3.7 Å respectively. Using our approach of iterative response prediction and kinetic excitations to overcome free energy barriers, we identified the most-stable ligand-bound conformations (predicted holo-structures) alongwith the transition intermediates. These predicted holo-structures matches closely with the experimental holo-structure as the RMSD between the two for the case of ADK, BGT and RBP was 2.30, 1.58 and 1.40 Å respectively. We defined an energy-based scoring function, which includes changes in MM/PBSA protein-ligand interaction energy and protein strain energy, to identify the most stable ligand-bound state. For all three proteins, among all predicted conformations, the most-stable ligand-bound state identified using the energy function is structurally closest to the experimental ligand-bound structure. Further, for the case of ADK and RBP, we compared the predicted transition pathway with the transition pathway obtained from bias-exchange metadynamics and umbrella sampling simulations respectively. In the respective CVs space of both cases, we observed a close agreement between the transition pathway predicted by our approach and those obtained from the low-throughput methods in the previous studies. Crystallographic structures of RBP provides an approximate view of the path of complete transition in the order, 1*BA2_A* → 1*BA*2_*B* → 1*URP* → 2*GX*6 → 2*DRI*. Our predicted intermediate states closely matches with the crystallographic proposed intermediates with the correct order of transitions. For example, intermediate *I*_1_ is closest to crystal structure 1*B*A2_*B* with rmsd of 1.39Å, intermediate *I*_2_ is closest to crystal structure *1URP* with rmsd of 1.73Å and final structure is closest to crystal structure 1URP with rmsd of 1.58Å. Also, for the case of RBP, among the intermediate states, we identified an intermediate which is possibly involved in ribose release. Interestingly, this intermediate is structurally closest to the intermediate state identified using umbrella sampling in a previous study.

To reach the final state, the method performs a series of small structure perturbations along the LRT-predicted direction. We have shown that, LRT-predicted response depends upon the conformational state of the protein and changes along the transition pathway as protein jumps from one intermediate state to another. Assuming residue fluctuations in a local free energy minima follows a Gaussian distribution, we derived that LRT-predicted initial response in the local minima is valid even for strong ligand-induced perturbation in protein Hamiltonian. For ADK, we observed a state-dependent dynamic coupling between the motions of LID and NMP domains which explains the characteristic non-linear transition pathway (LID closing followed by NMP closing) of ADK upon binding with its ligand Ap5A. In state *I*_1_, which is obtained after 1 *ns* MD relaxation of initial docked state, closure of LID domain was anti-correlated with opening of NMP domain whereas in intermediate state *I*_4_, closing of LID domain was correlated with the closing of NMP domain. Since, large scale protein transition pathways are typically non-linear, intermediate states plays an important role in modulating the structural response. We compared two approaches (i) a single large structure perturbation along the LRT-predicted direction, in which the structural response is predicted only initially and role of intermediate states is not accounted and (ii) series of multiple small structure perturbations along the LRT-predicted directions, in which the response is re-calculated after each perturbation. Holo-structures predicted using the first approach has shown lower similarity with the experimental structures as compared to the second approach. In first approach, the RMSD between predicted holo structures and crystallographic holo structures for AdK, RBP and BGT was 3.79, 3.94 and 2.36 respectively. Moreover, protein structure improvement, estimated as decrease in RMSD with the experimental holo structure after excitation, has strong correlation (0.91, 0.84 and 0.96 for ADK, RBP and BGT) with structure displacement along LRT-predicted direction for small displacements. The correlation start to decrease for higher displacements and even becomes negative for very high displacement values. This suggests that LRT-predicted response is most accurate to predict small-scale response and LRT-predicted structural response should be updated along the pathway. Thus, it is important to perform small scale structure perturbations crossing only one or few energy barriers in a single step, wherein after each structure perturbation, the changes in residue fluctuations and fluctuation co-variances due to change in conformational state are incorporated.

Perhaps, the most important feature of this protocol is that, it does not require any knowledge on the end state or the reaction coordinate to predict the ligand-induced protein structure transitions. Dominant normal modes triggered by ligand binding are predicted at each intermediate steps along the transition pathway and used as CVs for structure perturbation. Although the protocol doesn’t provide a continuous transition pathway but a coarse information on the characteristics of transition path is revealed by the predicted intermediate states. Interestingly, the dominantly triggered normal modes predicted at each step could be used as CVs in enhanced sampling method such as Metadynamics to gain accurate information of the transition free energy landscape. Also, the intermediate states predicted by our method could be used as inputs in path-sampling methods to obtain kinetics and thermodynamics of ligand-binding.

## Methods

### Protein-ligand complex preparation

Unbound protein structures are obtained from PDB database (AdK: *4AKE*, BGT: 1*JEJ* and RBP: 1*BA*2_*A*). Ligand 3D structures are taken from crystallographic holo structures of proteins in PDB database (AdK: 1AKE, BGT: 1JG6 and RBP: 2*DRI*) and missing hydrogen atoms are added using Avogadro.^48^. Missing hydrogen atoms in apo structure are added using the *pdb2gmx* tool of GROMACS (Version 5.1.4).^49^ Ligand is docked onto the protein structure binding site using software Hex (Version 8.0.0)^50^ and Autodock Vina^51^. Coordinates of ligand centre of mass obtained after superimposing the holo structure on apo structure are taken as reference ligand site. During ligand docking, ligand and protein structure are kept rigid and ligand translation search is constrained within a cubic box of side 40 Å centered at reference ligand site. Top 100 protein-ligand (PL) complexes, based on scoring function, are solvated with TIP3P water model^52^ and energy minimized using conjugate-gradient algorithm implemented in GROMACS^49^. Ligand parameters are generated using the *Ligand Reader & Modeler* module^53^ of CHARMM-GUI^54^. After energy minimization, these 100 PL complexes are re-ranked on the basis of protein-ligand MM/PBSA55 binding energy and most-stable PL complex complex is selected. MM/PBSA binding energies are calculated using *g_mmpbsa*^56^ and detailed calculation parameters are provided in *SI Appendix*. Selected complex is equilibrated with position restraints for 100 ps under isothermal-isochoric conditions (NVT ensemble) and 100 ps under isothermal-isobaric conditions (NPT ensemble). At this stage, the system is well equilibrated at desired temperature and pressure conditions *(T =* 310 K and *P* = 1 *atm)* and then, we performed a 1 ns long explicit-solvent NPT production simulation to observe the allow small-scale adjustments in PL complex. Solvation, energy minimization and MD simulations are performed using using CHARMM36 all-atom additive forcefield^57^ implemented in GROMACS (Version 5.1.4)^49^ and detailed simulation parameters of each stage are provided in *SI Appendix.* Mean squared fluctuations of residue Cα positions is calculated from isotropic B-factors using equation 14, which are obtained using *rms f* module of GROMACS on a 1 *ns* trajectory of apo protein in solution.

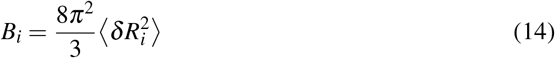

### Normal modes, residue fluctuations and covariance matrix calculation

Normal modes of a protein (*N* residues) in a given conformation **X_0_** are calculated by representing it as an anisotropic network model (ANM)^58^, in which each residue is modelled as a single bead centered at the position of its C-alpha atom and each bead in the model is connected to all other beads located within a cutoff radius *R_c_* by harmonic springs of uniform spring constant *γ*. Selection of ANM parameters *(R_c_, γ*) is discussed in Results section. Normal modes are obtained by eigendecomposition of mass-weighted Hessian matrix 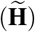.^58^

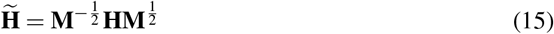

**M** is 3*N* × 3*N* diagonal mass matrix containing mass of each residue thrice, **H** is 3*N* × 3*N* Hessian matrix whose elements corresponds to the partial double derivatives of the potential energy function *E* (**X**) and, are given as

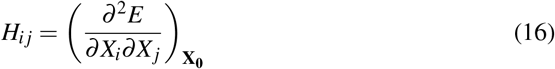

For a simplified representation, the conformation of a protein (N residues) is expressed in generalized coordinates **X** (3N dimensional coordinates of C-alpha atoms of residues) such that **X**^*T*^ = (*X*_1_,*X*_2_...*X*_3*N*_)^*T*^ = (*x*_1_,*y*_1_,*z*_1_,*x*_2_...*z_N_*)^*T*^. The potential energy function of ANM is given in equation 8. Eigenvectors (**U_i_**) of 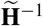 are the direction of normal modes and their corresponding eigenvalues (*λ_i_*) gives the fluctuation frequencies 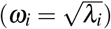 along these normal modes. First six eigenvectors (zero eigenvalues) which corresponds to the rotational and translational motion of protein are ignored. Covariance matrix of protein residue fluctuations is obtained as^58^

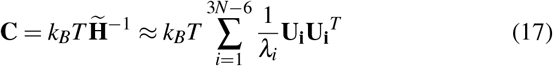

Diagonal elements of **C** corresponds to variance of residue fluctuations along each axis and thus mean fluctuations of residues is estimated using ANM as^58^

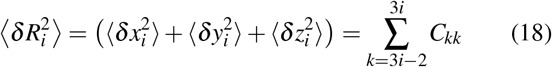

### Calculation of ligand-induced force on residues and LRT-predicted direction of conformational change

A model based on protein-ligand MM/PBSA^55^ binding energy calculations is developed to estimate the perturbative force applied by the ligand on each protein residue. Energy contribution of each residue to the total binding energy is calculated using the energy decomposition scheme of *g_mmpbsa*^56^ for frames of a short MD trajectory. For any given residue, its binding energy contribution varies with change in its spatial position relative to the ligand. Thus, for each residue *i*, a quadratic potential energy hyper-surface model is fitted which best models its variation in binding energy contribution 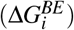 vs its spatial position relative to ligand (*x_i_, y_i_, z_i_*).

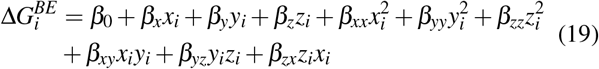

Ligand-induced perturbative force on *i^th^* residue is estimated as negative gradient of at mean residue position relative to the ligand during the trajectory.

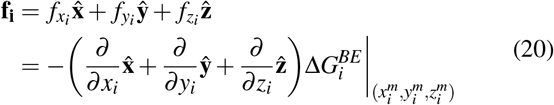

For simplification, LRT-predicted response of all residues (equation 2), is combined together in vector form, 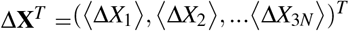, which is calculated as

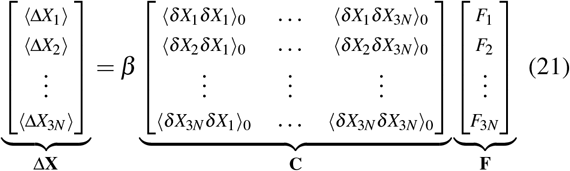

**C** is the covariance matrix of protein-residue fluctuations (equation 18) and *F_i_*s are the forces exerted by the ligand on residues such that {*F*_1_, *F*_2_,... *F*_3*N*_ } = {*f*_*x*_1__, *f*_*y*_1__,... *f_z_N__* }. Unit vector along Δ**X** is referred to as the LRT-predicted direction of conformational change (**d**_*LRT*_).

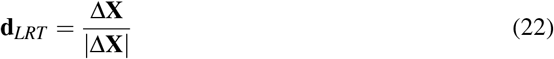

### Enhanced sampling along LRT-predicted direction and selection of output structures

Enhanced sampling along the LRT-predicted direction is performed using MDeNM. Protein motion is excited along the direction which is linear combination of relevant modes. Selection of mode coefficients in the linear combination is discussed in Results section. Protein structure change is observed at varying degree of excitation. Mean residue displacement (*D_P_*) of a protein (*N* residues) structure **X** from the initial structure **X_0_** along the excited direction (**d**_*LRT*_) is calculated as

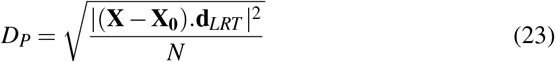

A three-point average of *D_P_* vs. Δ*T* data shows the presence of several palteaus which corresponds to the range of Δ*T* in which *D_P_* fluctuates around a constant value. Mathematically, plateaus corresponds to a region of low variance in average *D_P_*, thus we calculated a 3-point local standard deviation *(LSD)* of average *D_P_* vs Δ*T*. Local minimas in *LSD* vs Δ*T* data corresponds to the plateau regions and two adjacent plateaus are separated by a region of high variance which are identified by local maxima between the two minimas corresponding to the adjacent plateaus. Further details (with examples) are provided in *SI Appendix.* Structure corresponding to highest *DP* value in the plateau region is selected as the representative structure of the plateau.

### Structure similarity comparison and scoring analysis

3D structures of a protein, after superimposition, are compared using *RMSD*, which measures the root mean squared displacement of residue C*α* positions and *GDT_avg_*, which measures the average GDT (global distance test) score59 between two structure using cutoff values of 2, 4 and 8 Å. Lower the *RMSD* or higher the *GDTavg* between two structures, higher is the structure similarity. Mean residue displacement *(D_m_*) of a protein (*N* residues) structure **X** from the initial structure **X_0_** along the *m^th^* mode (**U_m_**) of initial structure is calculated as

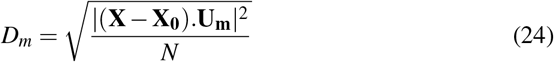

We defined a scoring function (ΔΔ*G*) to estimate the relative stability of predicted protein-ligand complexes compared to initial complex.

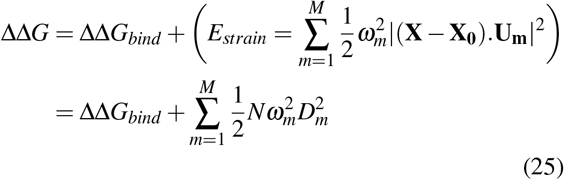

ΔΔ*G* accounts for changes in protein-ligand MM/PBSA binding energy (ΔΔ*G_bind_*) and protein strain energy *(E_strain_*) due to changes in protein conformation. *E_strain_* is estimated by assuming

### Implementation

Normal mode analysis, calculation of ligand-induced force on residues and LRT-predicted direction of conformational change are performed using MATLAB scripts.

## Supplementary information

### Prediction of direction of protein conformational change due to ligand binding

Let, a protein in solution environment is our system of interest, governed by its Hamiltonian 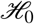. The partition function of the protein (*Z*(*β*)) is described by equation A1.

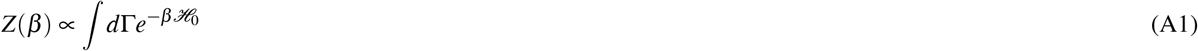

The integral is over phase space (Γ) and *β* = (*k*_B_*T*)^-1^ is the thermodynamic beta. Ligand binding perturbs the Hamiltonian of unbound-protein and protein change its state to reach the new free energy minima. The Hamiltonian of ligand-bound state 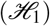 is described as perturbation to the Hamiltonian of ligand-unbound state 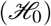 (equation A2), while considering the ligand as a part of the environment.

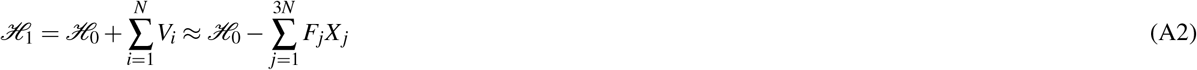

*V_i_* is the interaction energy between ligand and *i^th^* residue of protein, *N* is the number of protein amino-acid residues, *X_j_*s are the generalized coordinates such that {*X*_1_,*X*_2_,... *X*_3*N*_} = {*X*_1_,*y*_1_,... *z_N_*} and Fs are the forces exerted by the ligand on residues such that {*F*_1_,*F*_2_,... *F*_3*N*_x} = {*f*_*x*_1__, *f*_*y*_1__,... *f_Z_N__*x}, where {*x_i_,y_i_,z_i_*} are coordinates of *i^th^* residue and *f_x_i__*, *f_y_i__* and *f_z_i__* are the ligand-exerted forces on *i^th^* residue in x,y and z direction. The partition function of the perturbed protein (*Z*(*β*)) is described by equation A3.

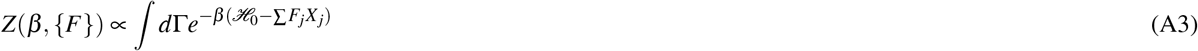

A large number of system properties can be derived from the natural log of partition function (ln (*Z*(*β*, {*F*}))). Thus, we write ln (*Z*(*β*, {*F*})) as a Taylor expansion with perturbation forces as variables (equation A4).

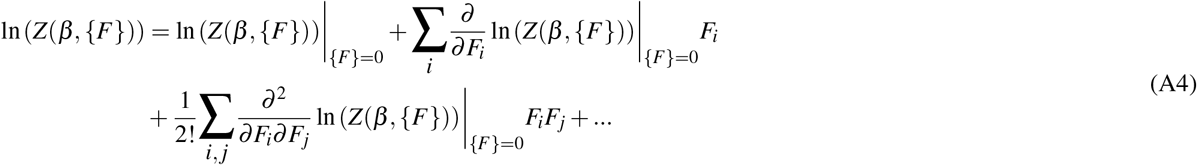

The first order partial derivatives in equation A4 can be derived by partial differentiation of equation A3 on both sides. The first order partial derivatives gives the ensemble averages of residue coordinates (equation A5).

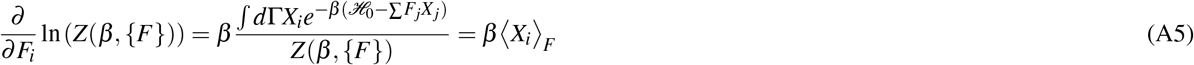

〈·〉_*F*_ represents the ensemble average of a system property in the perturbed state. Similarly, higher order derivatives in equation A4 can be derived from equation A3. For example, second order derivatives gives the co-variance of residue position coordinates (equation A6), third order derivative gives coskewness of residue position coordinates (equation A7) and in general, the *n^th^* order partial derivatives can be derived as equation A8.

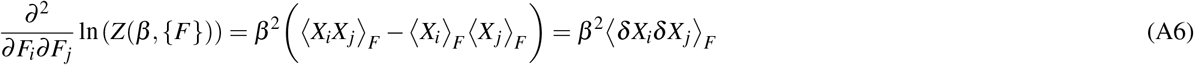

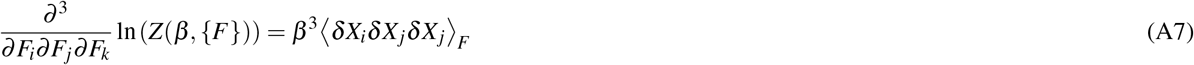

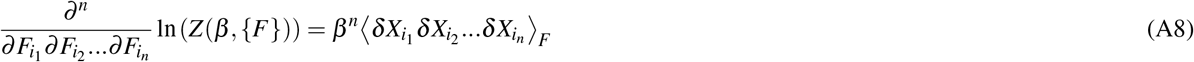

In equation A6, A7 and A8, *δX_i_* = *X_i_* — 〈*X_i_*〉. Since, our purpose is to predict the structural response of protein upon ligand binding, equation A5 suggests that the average coordinates of residue positions in the perturbed state can be obtained by first order partial derivatives of ln (*Z*(*β*, {*F*})) with respect to perturbation forces. Thus, taking a partial derivative of equation A4 with respect to *F_m_* and multiplying both sides by *β* gives

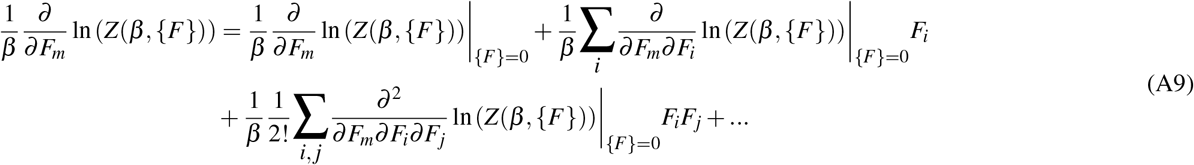

Using, equation A5–A8, equation A9 can be written as

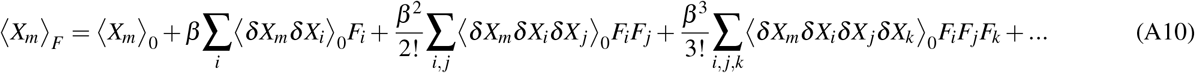

Equation A10 gives the ensemble average of residue position coordinates in the perturbed state in terms of the residue coordinates before perturbation, perturbation forces, residue fluctuations and fluctuation correlations before perturbation. This equation, mathematically, shows the role of intrinsic motions in governing the conformational change of protein.

### Special case 1: Weak perturbation

For a weak perturbation in protein Hamiltonian (*F_i_. = small ∀ i*), higher order terms in equation A10 can be ignored and the resulting equation is called as linear response.

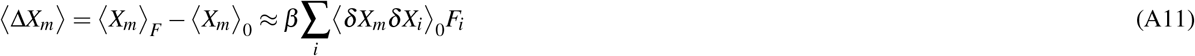

### Special case 2: Strong perturbation and Gaussian fluctuations

Assuming a harmonic approximation of potential energy of protein, which is valid for small fluctuations near minima of an energy well, residues displacement from its mean position follows a multivariate Gaussian distribution i.e. the variable *δ***X** = [*δX*_1_, *δX*_2_,..., *δX*_3*N*_] is normally distributed defined by mean **0** and covariance matrix **C**. For a variable with zero-mean multivariate Gaussian distribution, its higher order moments *m_n_* = 〈*δX*_*i*1_ *δX*_*i*2_...*δX_i_n__*〉 can be easily determined from the covariance matrix using Isserlis’s theorem (also known as Wick’s probability theorem).60 Using the theorem, when *n* is odd, the moment (*m_n_*) is zero and when *n* is even, the moment (*m_n_*) is given as

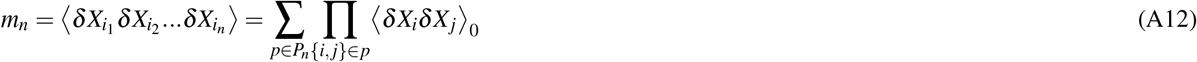

In equation A12, *P_n_* is a set comprising all different ways of partitioning {*i*_1_, *i*_2_,..., *i_n_*} into pairs {*i, j*}. The sum is over all different ways of partitioning and the product is over all the pairs in a partition *p*. Using equation A12, equation A10 can be written as

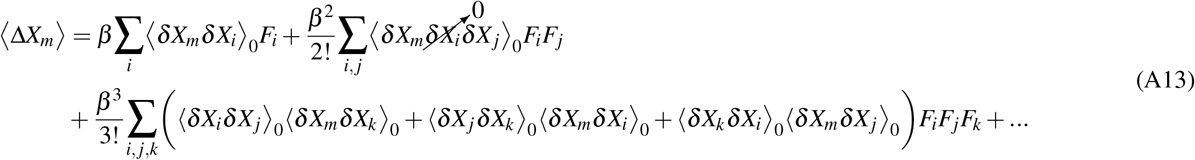

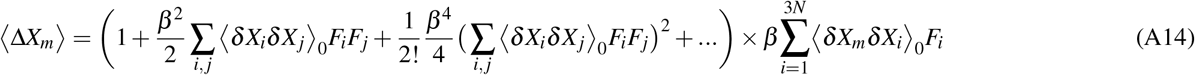

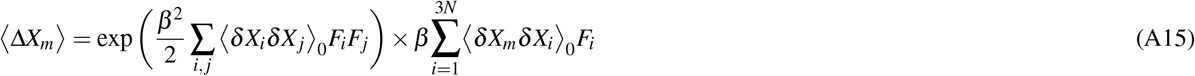

Equation A15 shows that the structural response of protein is proportional to the linear response under the assumption of Gaussian fluctuations of residue positions or, in other words, direction of structural response is same as predicted by the linear response formula (equation A11). This, implies that linear response formula (equation A11) can be used to predict the direction of initial structural response of protein even for strong ligand-induced perturbations.

## Supplementary Figures and Tables

**Figure S1.**
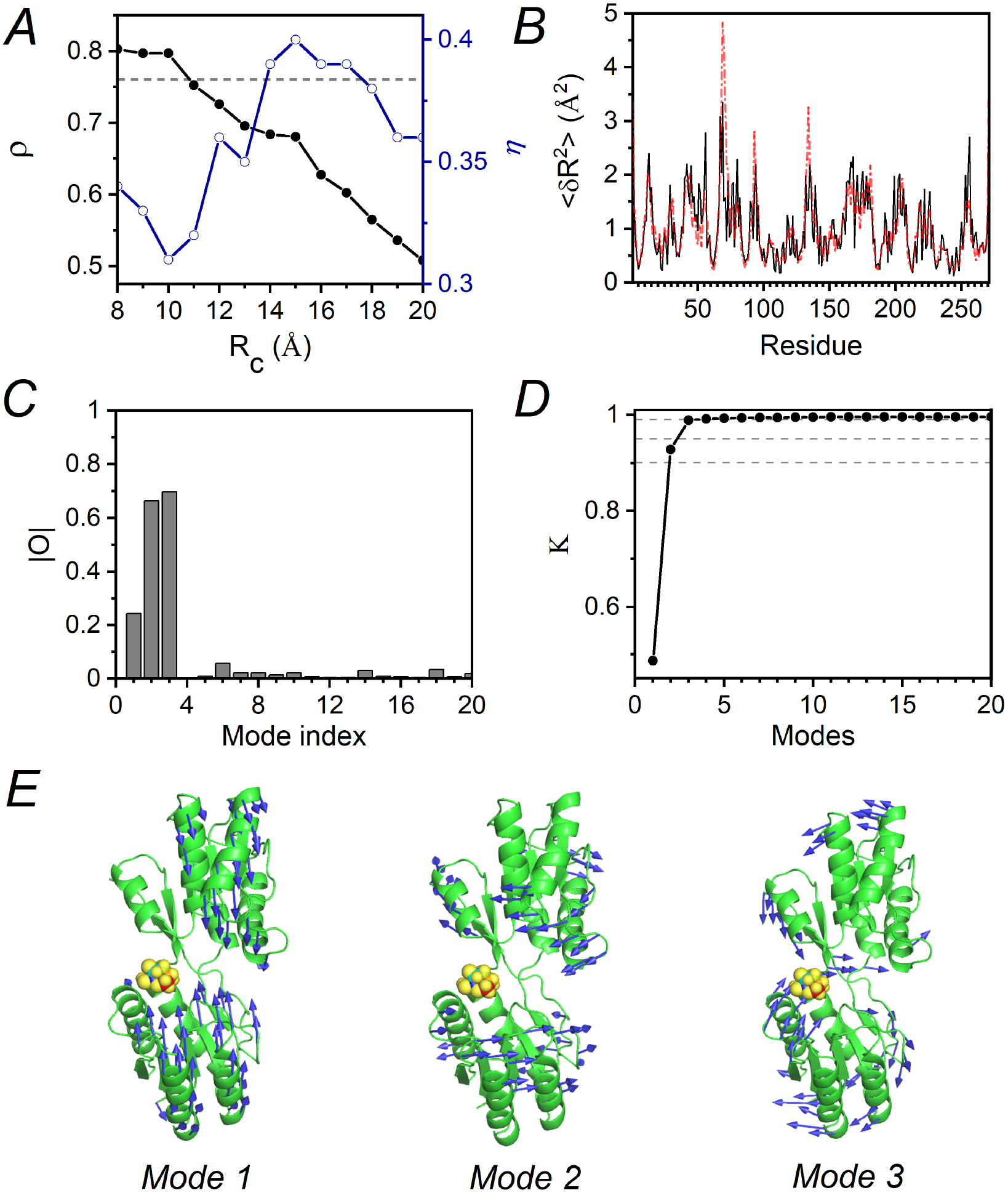
Estimation of ANM parameters and identification of normal modes triggered by ligand binding (Ribose Binding Protein). (**A**) Pearson correlation (*ρ*) between residue mean squared fluctuations predicted by ANM and estimated by 1 ns long MD simulations at different *R_c_* values (filled circles, left Y-axis). A cutoff line at *ρ* = 0.95 × *ρ_max_* is shown (dashed line). Mode contribution diversity estimated by η for different *R_c_* values (empty circles, right Y-axis). (**B**) Mean squared fluctuations of residues in apo-state obtained from 1ns MD simulation (dotted line) and estimated by ANM at *R_c_* = 10Å (solid line). (**C**) Overlap (absolute value) of first 20 normal modes with LRT-predicted direction. (**D**) Cumulative contribution of top normal modes to LRT-predicted direction in decreasing order of overlap (circles) and reference lines at *K* = 0.9, 0.95 and 0.99 (dotted lines). (**E**) Direction of motion of residues in *Mode* 1, *Mode* 2 and *Mode* 3 shown by blue arrows on cartoon diagram of protein-ligand complex (ligand atoms are shown as spheres).

**Figure S2.**
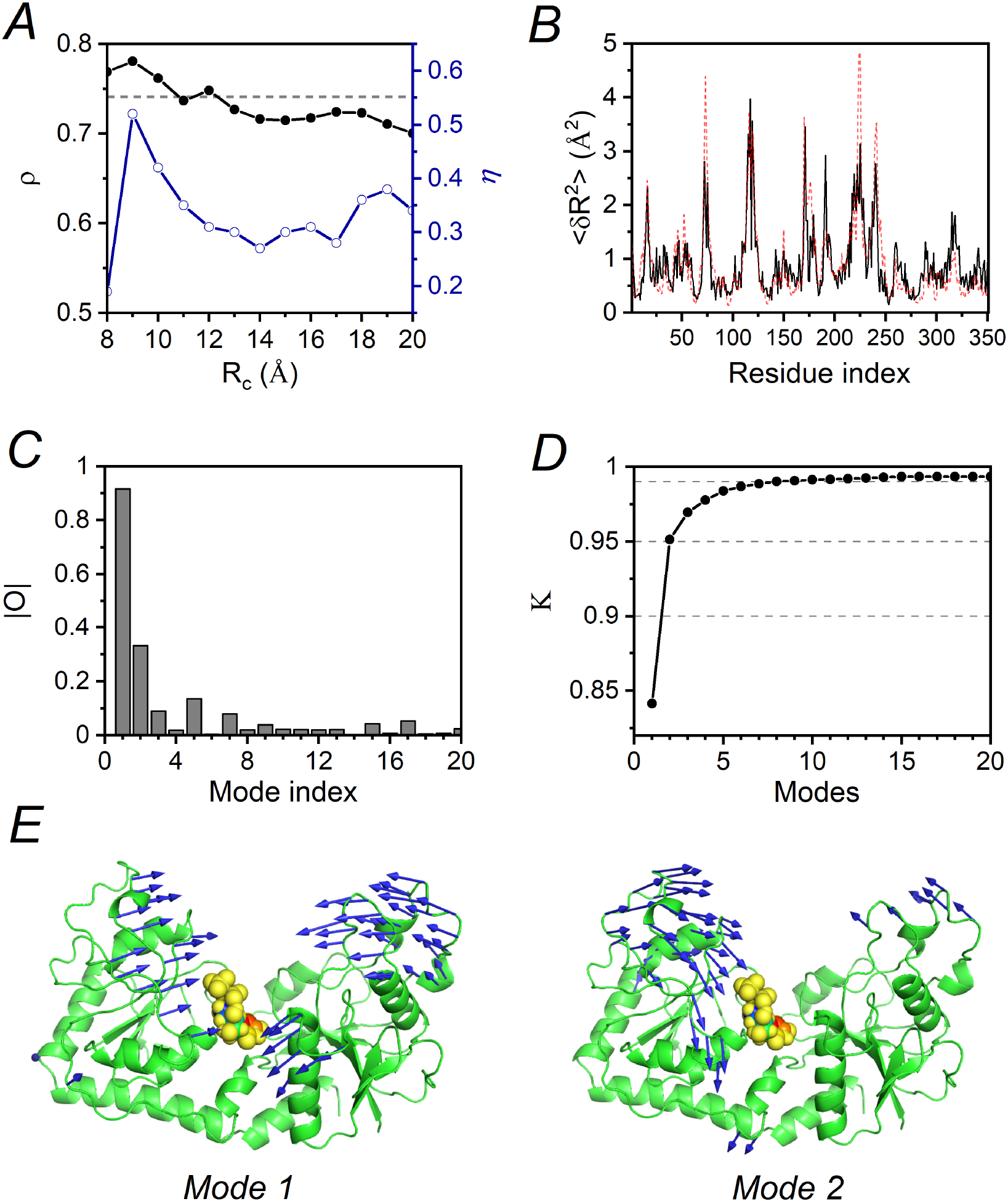
Estimation of ANM parameters and identification of normal modes triggered by ligand binding (*β*-glucosyltransferase). (**A**) Pearson correlation (*ρ*) between residue mean squared fluctuations predicted by ANM and estimated by 1 ns long MD simulations at different *R_c_* values (filled circles, left Y-axis). A cutoff line at ρ = 0.95 × *ρ_max_* is shown (dashed line). Mode contribution diversity estimated by η for different *R_c_* values (empty circles, right Y-axis). (**B**) Mean squared fluctuations of residues in apo-state obtained from 1ns MD simulation (dotted line) and estimated by ANM at *R_c_* = 8Å (solid line). (**C**) Overlap (absolute value) of first 20 normal modes with LRT-predicted direction. (**D**) Cumulative contribution of top normal modes to LRT-predicted direction in decreasing order of overlap (circles) and reference lines at *K* = 0.9, 0.95 and 0.99 (dotted lines). (**E**) Direction of motion of residues in *Mode* 1 and *Mode* 2 shown by blue arrows on cartoon diagram of protein-ligand complex (ligand atoms are shown as spheres).

**Figure S3.**
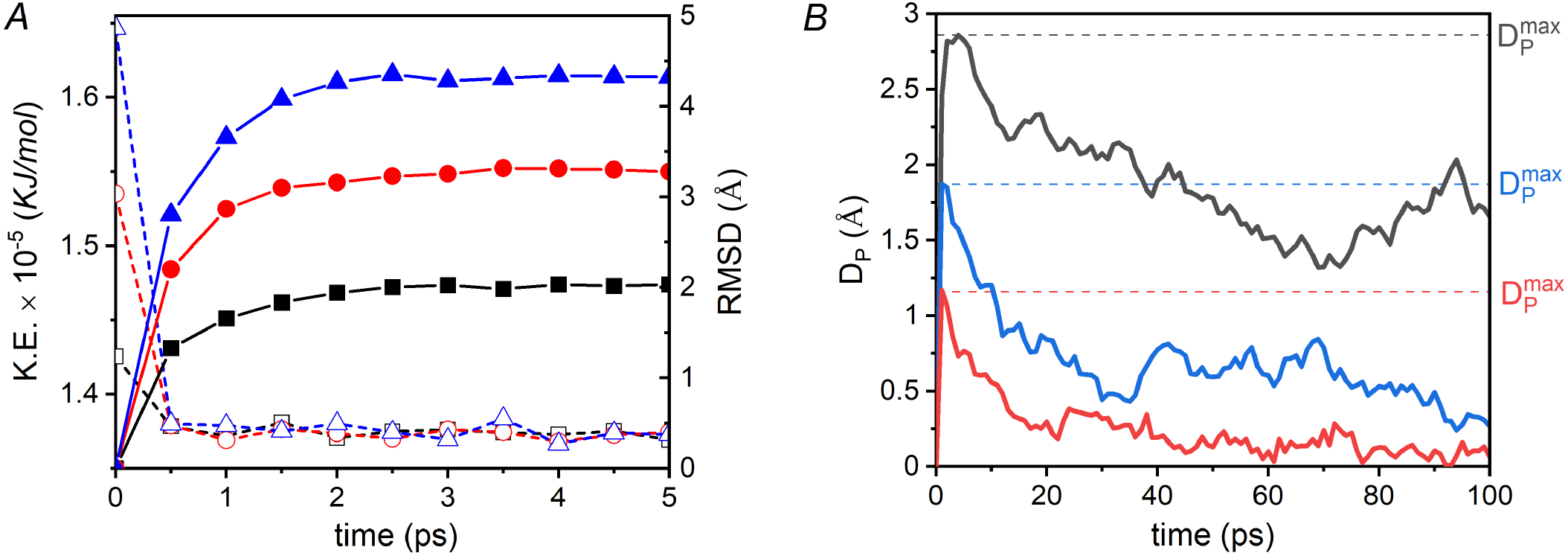
Molecular dynamics with excited normal modes (MDeNM). (**A**) Variation of kinetic energy of the AdK in water system (empty symbols) and *RMSD* of protein from initial structure (filled symbols) in a MDeNM excitation of 5 *ps* along *Mode* 1 at excitation temperature Δ*T* =10*K* (black), 30*K* (red) and 50*K* (blue). (**B**) Displacement along the excited direction (*D_P_*) in a 100 ps MDeNM excitation of AdK along *Mode* 1, *Mode* 5 and *Mode* 9 at excitation temperature Δ*T* = 25 *K*. Maximum displacement along the excited direction 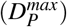 are marked for each case with dashed lines.

**Figure S4.**
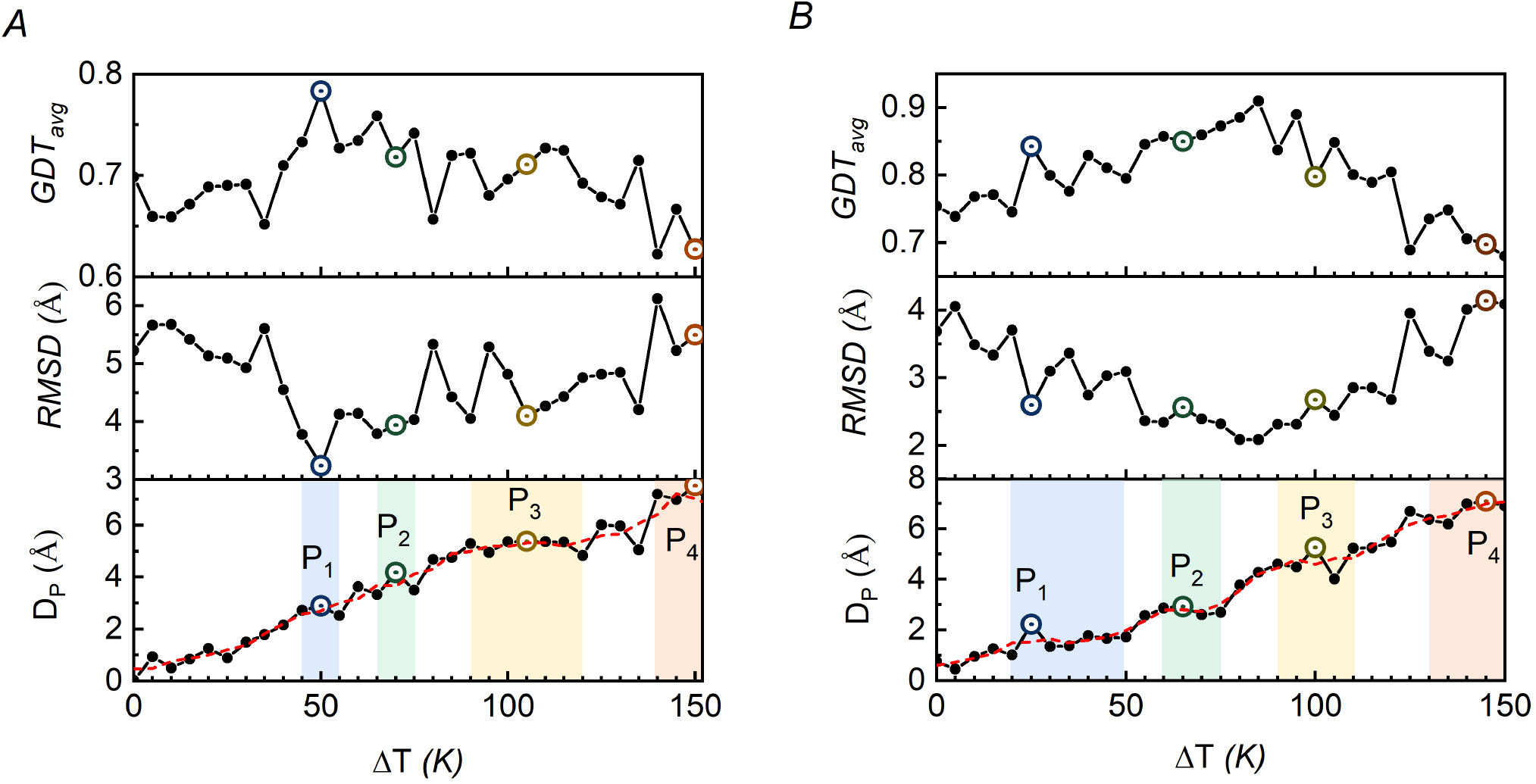
Enhanced sampling along LRT-predicted direction. Structure displacement along LRT-predicted direction (*D_P_*) in 100 ps MDeNM excitations at varying Δ*T* (circles, solid line) along with three-point average of *D_P_* vs. Δ*T* data (dotted line). Identified plateau regions in average *D_P_* vs. Δ*T* plot are highlighted (bottom panel). *RMSD* (middle panel) and *GDT_avg_* (top panel) score between the experimental holo structure and structure obtained at the end of MDeNM excitation. Representative structure of each plateau is highlighted (all panels) for (**A**) Ribose Binding Protein and (**B**) *β* -glucosyltransferase

**Figure S5.**
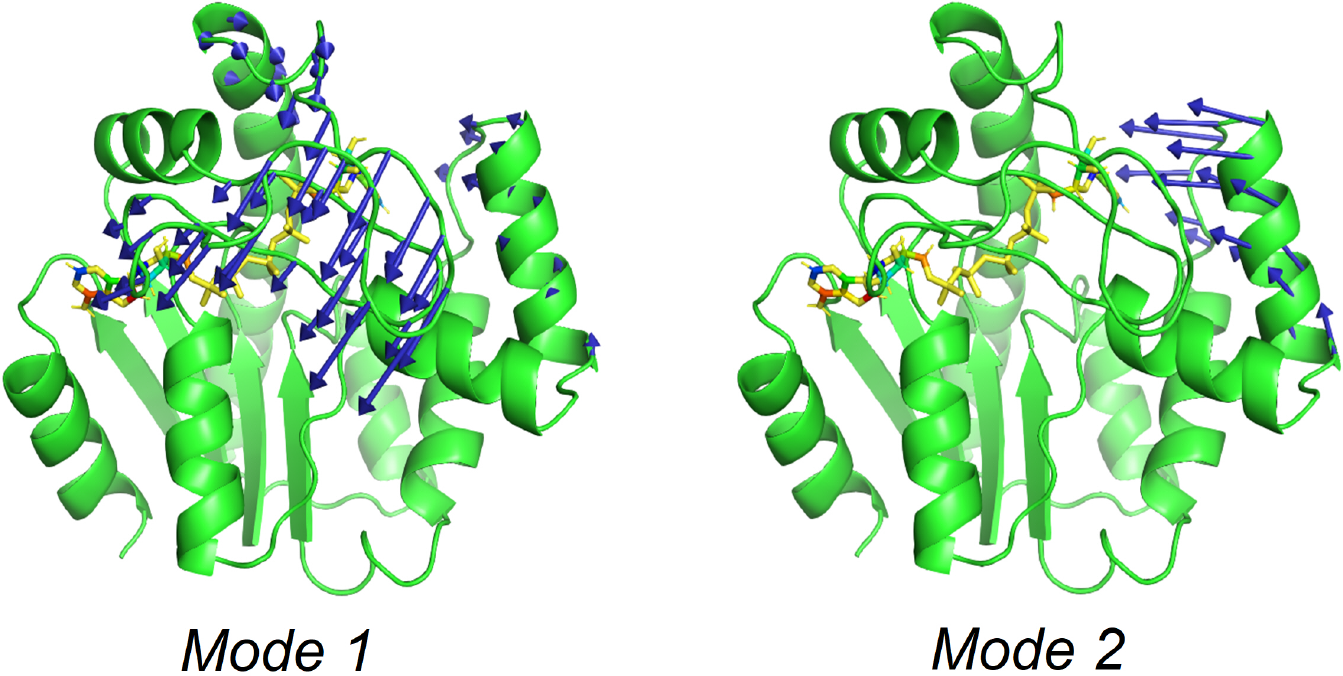
Dominant normal modes triggered by ligand in the intermediate state *I*_4_ of AdK. Structure cartoon diagram (green), ligand (yellow sticks) and residue direction of motion (blue arrows) are shown for *Mode* 1 and *Mode* 2 of intermediate state *I*_4_ obtain by our protocol during apo to holo transition of AdK

**Figure S6.**
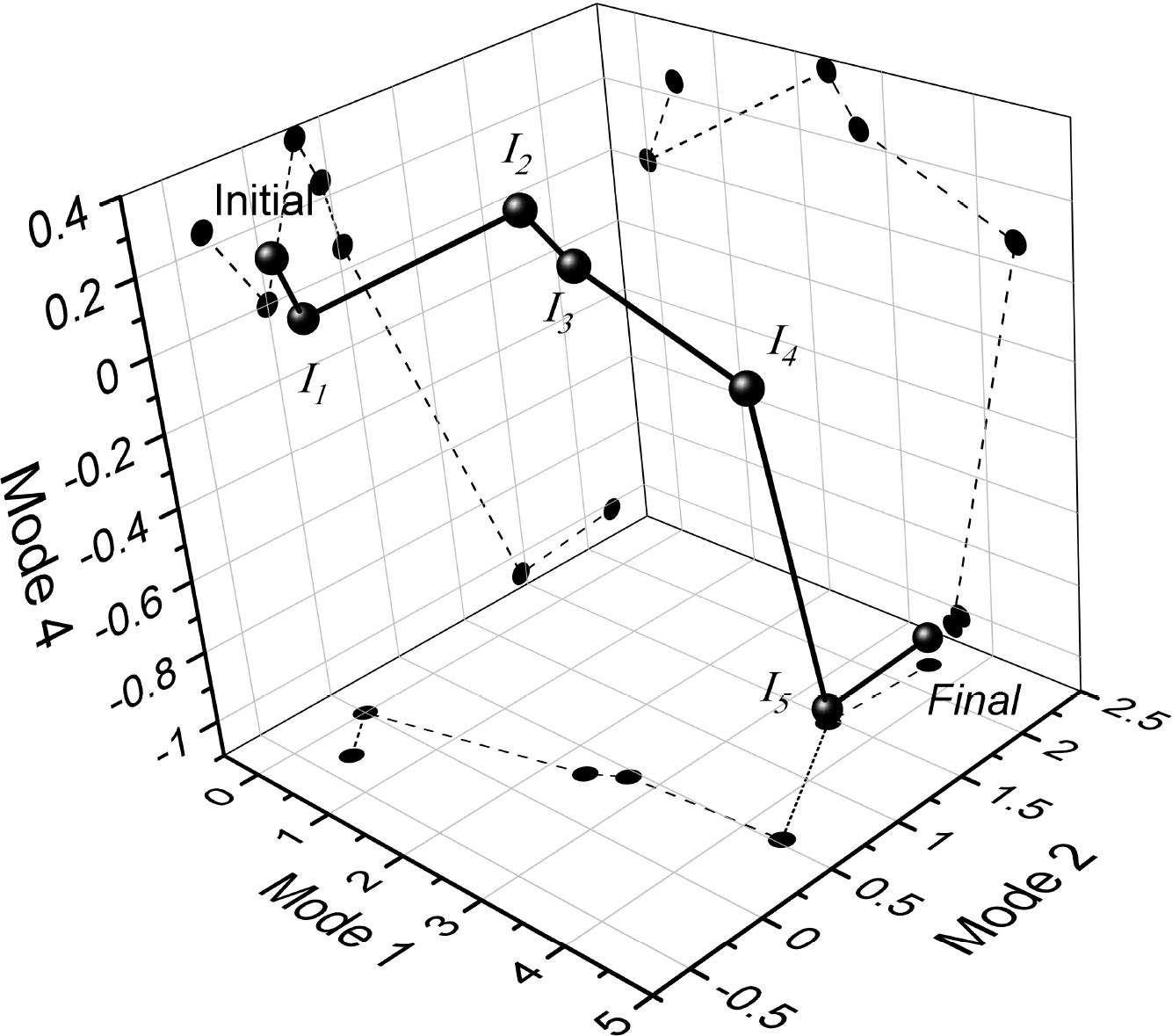
Adenylate Kinase transition pathway in normal mode space

**Table S1.** Comparison of actual and LRT-predicted magnitude of structural response

| Protein | AdK | RBP | BGT |
| --- | --- | --- | --- |
| Actual $RMSD$ (Å) | 7.1 | 6.2 | 2.1 |
| LRT-predicted $RMSD$ (Å) | 17.35 | 4.9 | 5.46 |

**Table S2.** Correlation between *D_P_* (displacement along LRT-predicted direction) and structure improvement (measured as decrease in *RMSD* with experimental holo structure)

| $\Delta T$ range | AdK | RBP | BGT |
| --- | --- | --- | --- |
| $0 - T_1^{end}$ | 0.91 | 0.84 | 0.96 |
| $T_1^{end} - T_2^{end}$ | 0.68 | 0.31 | 0.85 |
| $T_2^{end} - T_3^{end}$ | 0.26 | -0.05 | -0.60 |
| $T_3^{end} - T_4^{end}$ | -0.23 | -0.83 | -0.96 |
| $T_4^{end} - T_5^{end}$ | -0.70 | NA | NA |

$T_i^{end}$ = end of $i^{th}$ plateau NA = Not applicable

**Table S3.** Identification of holo struture from energetics (Ribose Binding Protein).

| | $RMSD(A^0)$ | $GDT_{avg}$ | $\Delta\Delta G(Kcal/mol)$ |
| --- | --- | --- | --- |
| *Initial* | 6.2 | 0.64 | - |
| $I_1$ | 5.23 | 0.69 | 0 |
| Single large perturbation |  |  |  |
| $P_1$ | 3.24 | 0.78 | -0.57 |
| **$P_2$** | **3.94** | **0.72** | **-0.65** |
| $P_3$ | 4.10 | 0.71 | +0.63 |
| $P_4$ | 5.49 | 0.62 | +2.00 |
| Multiple small perturbations |  |  |  |
| $E_1$ | 3.24 | 0.78 | -0.57 |
| $E_2$ | 2.09 | 0.88 | -1.44 |
| **$E_3$** | **1.58** | **0.92** | **-3.15** |
| $E_4$ | 1.90 | 0.91 | -0.98 |
| $E_5$ | 2.42 | 0.87 | -0.61 |
| $E_6$ | 3.01 | 0.80 | -0.28 |

**Table S4.** Identification of holo struture from energetics (*β*-glucosyltransferase).

| | $RMSD(A^0)$ | $GDT_{avg}$ | $\Delta\Delta G(Kcal/mol)$ |
| --- | --- | --- | --- |
| *Initial* | 2.1 | 0.90 | - |
| $I_1$ | 3.68 | 0.75 | 0 |
| Single large perturbation |  |  |  |
| $P_1$ | 2.59 | 0.84 | -3.15 |
| **$P_2$** | **2.36** | **0.84** | **-15.26** |
| $P_3$ | 2.67 | 0.79 | -6.72 |
| $P_4$ | 4.14 | 0.69 | -0.74 |
| Multiple small perturbations |  |  |  |
| $E_1$ | 2.59 | 0.84 | -3.15 |
| **$E_2$** | **1.40** | **0.96** | **-36.50** |
| $E_3$ | 1.49 | 0.95 | -21.67 |
| $E_4$ | 3.04 | 0.80 | -18.9 |
| $E_5$ | 2.94 | 0.84 | -3.43 |

